# Improved sequence mapping using a complete reference genome and lift-over

**DOI:** 10.1101/2022.04.27.489683

**Authors:** Nae-Chyun Chen, Luis F Paulin, Fritz J Sedlazeck, Sergey Koren, Adam M Phillippy, Ben Langmead

**Affiliations:** Department of Computer Science, Johns Hopkins University, Baltimore, MD, 21218, USA; Human Genome Sequencing Center, Baylor College of Medicine, Houston, TX, 77030, USA; Department of Computer Science, Rice University, 6100 Main Street, Houston, TX, USA; Genome Informatics Section, Computational and Statistical Genomics Branch, National Human Genome Research Institute, National Institutes of Health, Bethesda, MD, 20894, USA

## Abstract

Complete, telomere-to-telomere genome assemblies promise improved analyses and the discovery of new variants, but many essential genomic resources remain associated with older reference genomes. Thus, there is a need to translate genomic features and read alignments between references. Here we describe a new method called levioSAM2 that accounts for reference changes and performs fast and accurate lift-over between assemblies using a whole-genome map. In addition to enabling the use of multiple references, we demonstrate that aligning reads to a high-quality reference (e.g. T2T-CHM13) and lifting to an older reference (e.g. GRCh38) actually improves the accuracy of the resulting variant calls on the old reference. By leveraging the quality improvements of T2T-CHM13, levioSAM2 reduces small-variant calling errors by 11.4-39.5% compared to GRC-based mapping using real Illumina datasets. LevioSAM2 also improves long-read-based structural variant calling and reduces errors from 3.8-11.8% for a PacBio HiFi dataset. Performance is especially improved for a set of complex medically-relevant genes, where the GRC references are lower quality. The software is available at https://github.com/milkschen/leviosam2 under the MIT license.

## 1 Introduction

A reference genome serves both as a template for mapping reads and a set of coordinates for interpreting results. Human reference quality has steadily improved in the last decade, with over 1,000 GRCh37 issues having been resolved in GRCh38^1^, and T2T-CHM13 providing a telomere-to-telomere sequence including difficult-to-assemble regions like segmental duplications^2^.

While these improvements can benefit read mapping and downstream analyses^1, 3, 4^, researchers face obstacles when migrating between references. A new reference’s coordinate system is incompatible with annotations expressed in the old coordinates. Such annotations might be genotypes^5–7^, or functional or phenotype annotations^8–11^. Migrating from GRCh37 to GRCh38 required years of work^12^; even today, many groups report that they have no plans to switch from GRCh37^13^.

To facilitate movement between references, tools that “lift” across genomic coordinate systems have been proposed^14–17^. Unfortunately, the lifting process can produce discordant results compared to re-analyzing the sequencing reads from scratch. The problem is particularly pronounced in regions where the references have copy-number differences or assembly artifacts (Figure 1a)^12, 18–20^. While the recently described LiftOff method can lift gene annotations with high confidence using re-mapping^21^, this strategy works only with genes and not with other types of annotations such as generic intervals, genotypes, or read mappings.

**Figure 1:**
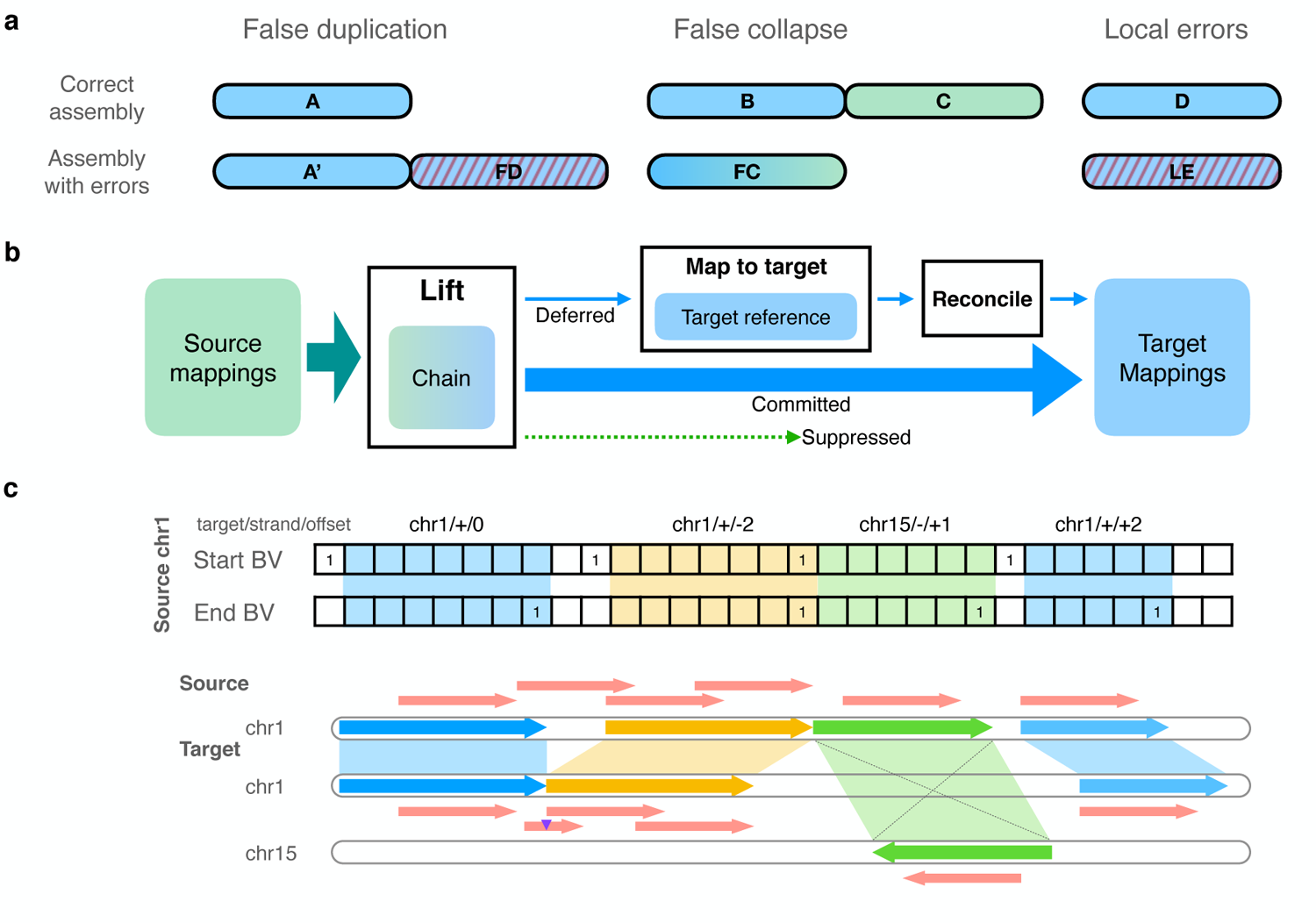
Overview of levioSAM2. **a**, Common assembly errors in legacy reference genomes include false duplications (FD), false collapses (FC), and local errors (LE). **b**, The levioSAM2 workflow; A chain file provides information to lift mappings. Mappings which cannot be confidently lifted are deferred and re-mapped. The final output is a set of mappings to the target reference after reconciling the deferred and committed mappings. Reads originating in duplications and local errors can benefit from the “commit” and “defer–reconcile” strategies, where mappings to the source are considered. False collapses can be resolved by the “suppress” strategy, which avoids spurious alignments in the collapsed reference by only including mappings from a single orthologous copy. **c**, LevioSAM2-lift uses a pair of succinct bit vector data structures to efficiently query chain segments. LevioSAM2 lifts aligned reads from source reference to target reference using a chain file and updates the alignment CIGAR. Blue, yellow, and green arrows represent chain segments. Red lines represent mapped sequences. The purple triangle represents a 2-bp insertion in the source reference which requires a CIGAR update.

The prior levioSAM software could perform scalable and memory-efficient lift-over of mappings, but did not support complex genomic rearrangements such as translocations and inversions^17^. Other methods either cannot lift read mappings^14, 16^ or do not scale for large genomic datasets^15^.

To make the best use of improved reference assemblies and rich annotations, we propose levioSAM2 for fast and accurate lift-over of read mappings (Figure 1b). In contrast to the more typical strategy of lifting old-reference alignments to a newer reference (old-to-new), we propose to start by mapping to the most complete and error-free assembly like T2T-CHM13, then to use levioSAM2 to lift the mapped reads to an existing annotation-rich reference like GRCh37 or GRCh38. The first step of levioSAM2 is an efficient lift kernel (“levioSAM2-lift”) that translates mapping information into the target coordinates. LevioSAM2-lift uses succinct data structures that can update mapped reference name and position information in *O*(1) time and update the CIGAR alignment string in *O*(*r*+*g*) time, where *r* is the number of CIGAR runs and *g* is the number of overlapping chain gaps of a mapping (Figure 1c and Section 4.1).

We further designed a selective strategy, similar to the “reference flow” approach^22^, to handle mappings that are influenced by major differences between source and target (Section 4.2). LevioSAM2 classifies lifted reads into three groups. The “suppressed” group consists of reads mapped to regions in the source genome with no counterpart in the target, e.g. the centromeric sequences in T2T-CHM13 and missing collapsed sequences, to avoid false-positive mapping. The “deferred” group consists of reads that can benefit from re-mapping, e.g. because they mapped with low mapping quality. The “committed” group, which generally contains the majority of the reads, consists of reads belonging to neither of the other two groups. Committed reads are mapped with high confidence and are included in the final mapping results (See Section 4.2). LevioSAM2 also collects and reports mappings that are unliftable using the coordinate system of the source reference, enabling analysis in regions unique to the source (Figure S1).

We evaluated levioSAM2 by lifting mappings from T2T-CHM13 to GRCh37 and GRCh38 coordinates and comparing them to the results obtained by mapping directly to the GRC references. We also called small variants using GATK-HaplotypeCaller^23^ and DeepVariant^24^. We observed that levioSAM2 could reduce small variant errors by 11.4% to 39.5% on real Illumina whole-genome-sequencing (WGS) data for HG001, HG002, and HG005 in the Genome in a Bottle (GIAB) v4.2.1 regions^25^. LevioSAM2 yielded larger improvements in the GIAB challenging medically relevant genes (CMRG)^26^, reducing small variant errors by 19.4% to 51.3% for HG002. LevioSAM2 also improved mapping of real Pacific Biosciences High Fidelity (PacBio HiFi) reads^27^. Besides achieving improved or comparable small-variant calling accuracy, levioSAM2 reduced structural-variant (SV) errors by 11.8% compared to GRCh38 in the GIAB CMRG regions^26^, and reduced SV errors by 3.8% compared to GRCh37 in the GRCh37 GIAB Tier 1 benchmark regions^28^.

## 2 Results

### Lifting from T2T-CHM13 improves short-read mapping to GRC references

We used simulated reads to compare mapping accuracy between the levioSAM2 workflow and a typical single-reference direct mapping method (Section 4.4). We can measure the correctness of a read mapping by comparing its mapping position with its true point of origin according to the simulator. We simulated 10M paired-end 100-bp Illumina reads using mason2^29^. We selected GRCh38 as the “base” reference, where sequences not liftable to CHM13 were masked with the `N` (unknown) symbol during simulation. We also injected the SNPs of HG001 during simulation (see Section 4.4 for details). To assess the influence of the quality of the reference assembly on the mapping process, we performed the simulation using both human chromosome 20 (GRCh38 chr20) and 21 (GRCh38 chr21). Chr21 is known to include 771 kbp of false duplications^26^, whereas chr20 has no known false duplications. False duplicates in the reference assembly can “attract” reads from the correct point of origin. We mapped the reads using Bowtie 2^30^ and BWA-MEM^31^, using default options for both.

The selective levioSAM2 workflow generated an additional 0.08% of correct mappings for chr20 and 1.37% for chr21 versus the direct-to-GRCh38 method (“GRCh38”; Figure S2). The fact that levioSAM2 had a more substantial improvement in chr21 reflects its ability to recover from the false duplicates in GRCh38. Note that since this simulation is based on GRCh38, we are not able to measure whether and how levioSAM2 benefits from assembly improvements in places where GRCh38 has assembly gaps, but characterize these improvements using real data in the following sections.

### LevioSAM2 improves short-read small variant calling

We next evaluated levioSAM2 using two common short-read small variant calling pipelines, BWA-MEM–GATK-HaplotypeCaller^23^ and BWA-MEM–DeepVariant^24^ (Section 4.5.1). We used real 30x paired-end Illumina Novaseq whole-genome sequencing (WGS) data representing three ancestries (HG001: Utah/European; HG002: Ashkenazi Jewish; HG005: Han Chinese)^32^ (Table 1). All three individuals had high-quality genotypes provided by the Genome in a Bottle (GIAB) consortium for both GRCh37 and GRCh38^25^. The GATK-based pipelines that used levioSAM2 to lift reads from T2T-CHM13 to GRCh37 (“CHM13-to-GRCh37”) reduced mean small variant error compared to a direct-to-GRCh37 pipeline by 49,770 errors (39.5%). LevioSAM2 also showed an error reduction when lifting from T2T-CHM13 to GRCh38 (“CHM13-to-GRCh38”), avoiding 18,308 (23.9%) small variant errors. The overall *F*_1_ improved as well (Fig 2a and Table S2). We stratified the calls into SNPs and indels and assessed the precision and recall for levioSAM2 and direct-to-GRC pipelines (Fig 2b). LevioSAM2 had higher precision and recall for all samples and both variant types. For all samples, levioSAM2 reduced the mean false-positive error (indel and SNP combined) by 38,981 (52.2%) and 11,551 (24.9%) compared to GRCh37 and GRCh38, respectively (Table S2).

**Figure 2:**
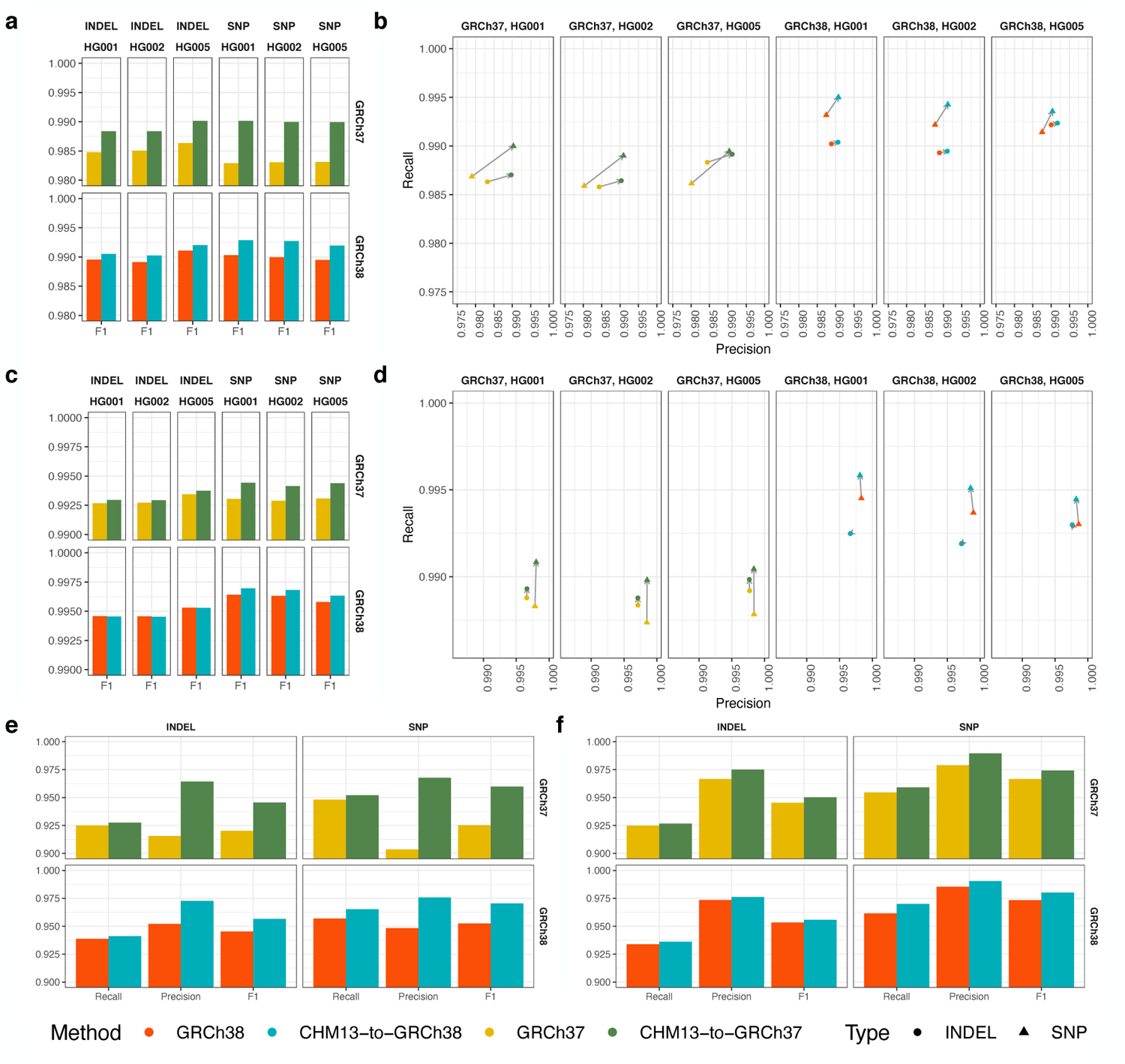
Small variant calling performance. **a, b**, Variant calling accuracy using GATK-HaplotypeCaller for all GIAB v4.2.1 regions (**a**: *F*_1_; **b**: recall and precision). **c, d**, Variant calling accuracy using DeepVariant for all GIAB v4.2.1 regions (**c**: *F*_1_; **d**: recall and precision). **e**, Accuracy for challenging medically relevant genes (CMRG) for HG002 using GATK-HaplotypeCaller. **f**, Accuracy for challenging medically relevant genes (CMRG) for HG002 using DeepVariant.

**Table 1:** Real sequencing datasets and truth sets we used for evaluation. The Illumina data are from Google Health^32^ and the PacBio data are from the Human Pangenome Reference Consortium^35^

| Sample | Sequencing platform | Variant type | Truth variants | Coordinates |
| --- | --- | --- | --- | --- |
| HG001 | 30× Illumina Novaseq | Small | GIAB v4.2.1<sup>25</sup> | GRCh37, GRCh38 |
| HG002 | 30× Illumina Novaseq | Small | GIAB v4.2.1<sup>25</sup> | GRCh37, GRCh38 |
| HG002 | 30× Illumina Novaseq | Small | GIAB CMRG<sup>26</sup> | GRCh37, GRCh38 |
| HG002 | 28× PacBio HiFi | Small | GIAB v4.2.1<sup>25</sup> | GRCh37, GRCh38 |
| HG002 | 28× PacBio HiFi | Small | GIAB CMRG<sup>26</sup> | GRCh37, GRCh38 |
| HG002 | 28× PacBio HiFi | SV | GIAB SV Tier 1<sup>28</sup> | GRCh37 |
| HG002 | 28× PacBio HiFi | SV | GIAB CMRG<sup>26</sup> | GRCh38 |
| HG005 | 30× Illumina Novaseq | Small | GIAB v4.2.1<sup>25</sup> | GRCh37, GRCh38 |

LevioSAM2 also improved small variant calling for the BWA-MEM–DeepVariant approaches. On average, the levioSAM2 workflows avoided 8,778 errors (16.7%) and 3,443 errors (11.4%) compared to GRCh37 and GRCh38, respectively (Fig 2c and Table S3). In contrast to the GATK-based pipelines, the strongest improvement using DeepVariant was for SNP recall, where levioSAM2 reduced mean false-negatives by 8,331 (20.8%) and 4,603 (22.3%) variants compared to GRCh37 and GRCh38 (Fig 2d and Table S3). The discrepancy between the performance of GATK-HaplotypeCaller and DeepVariant could be explained by the difference in their underlying models. The neural network model used by DeepVariant was capable of learning mapping artifacts and reducing false-positive errors in hard-to-map regions (Note S1). The hidden Markov model used by GATK-HaplotypeCaller might not have recognized the mapping artifacts, and thus benefited more from the less-biased mappings generated by levioSAM2.

Both GRCh37 and GRCh38 are known to include false duplications that confound genomic analysis in medically relevant regions. We compared levioSAM2, GRCh37, and GRCh38 for small variant calling performance in the GIAB CMRG regions^26^ using the HG002 WGS data. LevioSAM2-based methods were more accurate for both GATK-HaplotypeCaller and DeepVariant pipelines. When using GATK-HaplotypeCaller, the levioSAM2 workflows removed 1,514 (45.4%) and 730 (35.2%) variant calling errors compared to GRCh37 and GRCh38 (Figure 2e and Table S4). When using DeepVariant, levioSAM2 avoided 306 (19.4%) and 1,010 (20.0%) variant calling errors compared to GRCh37 and GRCh38 (Figure 2f and Table S5). The levioSAM2 workflows outperformed direct-to-target methods for both recall and precision for both variant types. LevioSAM2 had a larger improvement in the CMRG regions compared to other regions, again showing that levioSAM2 effectively leverages the improved assembly quality of T2T-CHM13 to improve short-read small variant calling.

We analyzed the reads mapped by CHM13-to-GRCh38 and called small variants in the “unliftable” regions that were unique to T2T-CHM13 using DeepVariant. We called 5,314 variants that passed the default DeepVariant filter, and 1,635 of them were high-quality (`QUAL` 30) (Figure S1). These variants were unique to T2T-CHM13 and could not be identified with typical variant calling approaches and GRCh38.

### Larger small variant calling improvement in difficult regions

We investigated differences in small variant calling between levioSAM2 and the GRC references using the GIAB difficult region stratifications^4, 26^. These stratify genomic regions by features such as low mappability (“LowMap“), extreme GC content (“GC*<*25or*>*65”), low complexity (“Tandem&Homo”), presence of segmental duplications (“SegDups”), and the assembly-artifacts-and-other regions (“OtherDifficult”). The union of these difficult regions comprises the “AllDifficult” regions (see Table S6 for region sizes). We considered only the variant calls within the GIAB confident regions in the evaluation. We observed that levioSAM2 had enriched small variant calling improvements in the LowMap, SegDups, and OtherDifficult regions when using GATK-HaplotypeCaller. In these regions, levioSAM2 had a 47.7%–48.0% and a 27.5%–42.4% reduction in error rate compared to direct-to-GRCh37 and direct-to-GRCh38 respectively. The union of all difficult regions also showed improved small variant calling performance (Figure 3a and Table S7). When using Deep-Variant, the most pronounced improvement was in LowMap regions, where levioSAM2 reduced errors by 15.7%–22.3% (Figure S5a and Table S8).

**Figure 3:**
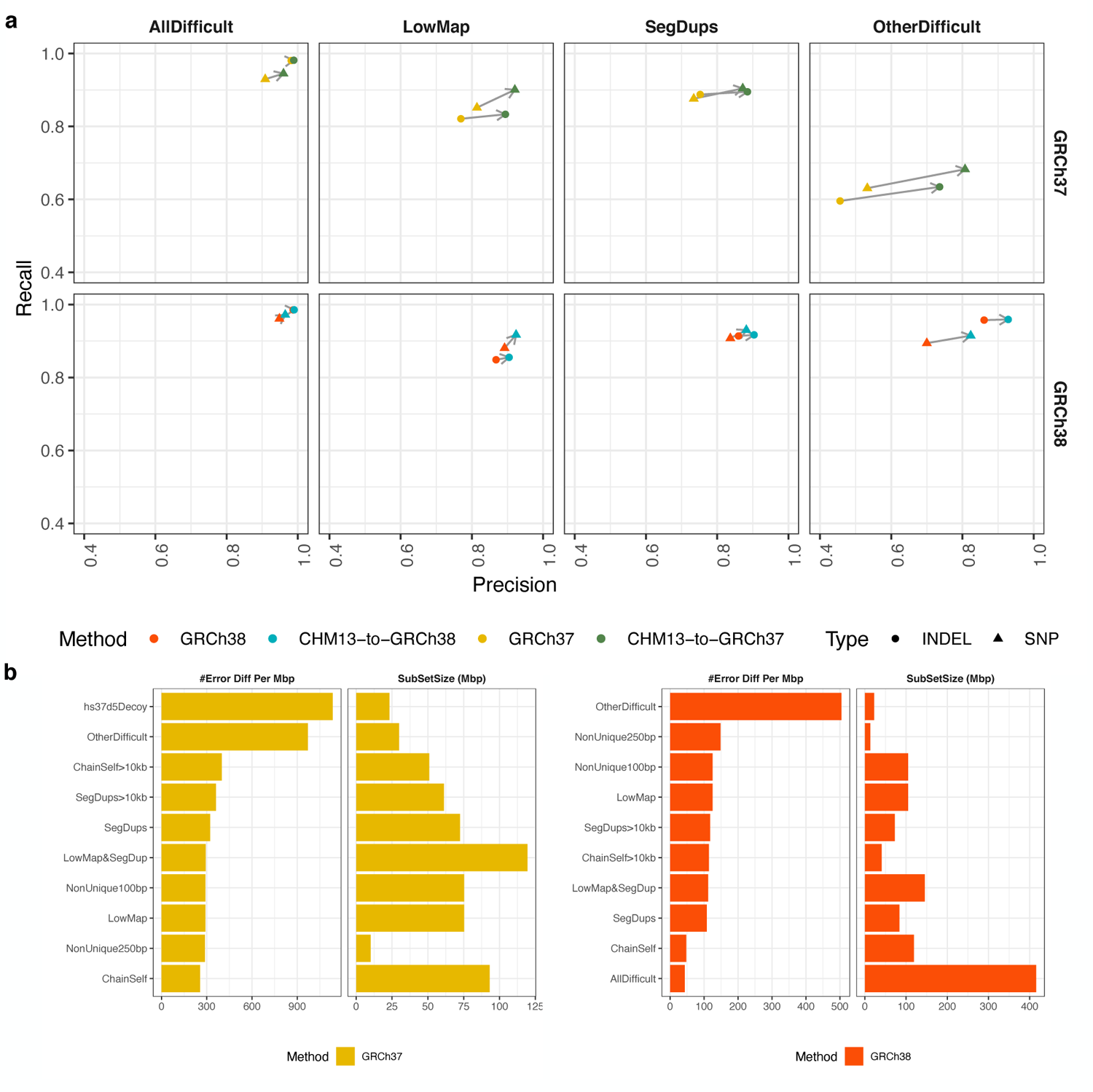
Small variant calling performance in difficult regions. **a**, Small variant calling accuracy in major difficult genomic regions for HG002. We excluded difficult regions with low complexity or extreme GC content in the plot because all methods perform similarly. **b**, GIAB stratified regions with top small variant calling error reduction densities by levioSAM2. Small variants in both plots were called using using GATK-HaplotypeCaller.

We further analyzed higher-resolution GIAB difficult region strata and ranked them by density of small variant calling error reduction (Figure 3b and Figure S5b). We observed that levioSAM2 reduced errors most in regions with likely reference artifacts (“hs37d5Decoy” and “ChainSelf”, both are subsets of “OtherDifficult”), avoiding up to 971 errors per Mbp compared to direct-to-GRC approaches. Other strata strongly improved by levioSAM2 included regions with large segmental duplications (“SegDups*>*10kb”), and low-mappability regions comprised of non-unique 250-mers (“NonUnique250bp”) and non-unique 100-mers (“NonUnique100bp”). There were no stratified regions that reported an increased error density of greater than 1/Mbp by levioSAM2 using either GATK-HaplotypeCaller or DeepVariant.

### Improved variant calling using PacBio-HiFi long reads

LevioSAM2 supports lifting alignments spanning chain gaps (Figure 1c), making it suitable for long reads, such as PacBio HiFi reads^27^. We designed a workflow for long reads supporting both minimap2^33^ and Winnowmap2^34^ (Section 4.2). We mapped a real 28× PacBio-HiFi WGS dataset from HG002^35^ to T2T-CHM13 and used levioSAM2 to generate GRC-based mappings. We called small variants using DeepVariant and assessed the calls with the GIAB v4.2.1 truth set. CHM13-to-GRCh37 removed 10,435 small variant errors (9.7%) compared to direct-to-GRCh37. CHM13-to-GRCh38 performed comparably to GRCh38, with 673 (0.8%) more errors (Figure 4a and Table S9). The levioSAM2 workflow generated more accurate small variant calls in the GIAB CMRG regions, where CHM13-to-GRCh37 avoided 30 (1.7%) errors and CHM13-to-GRCh38 removed 337 (22.5%) errors compared to their direct-to-GRC counterparts (Figure 4b and Table S10).

**Figure 4:**
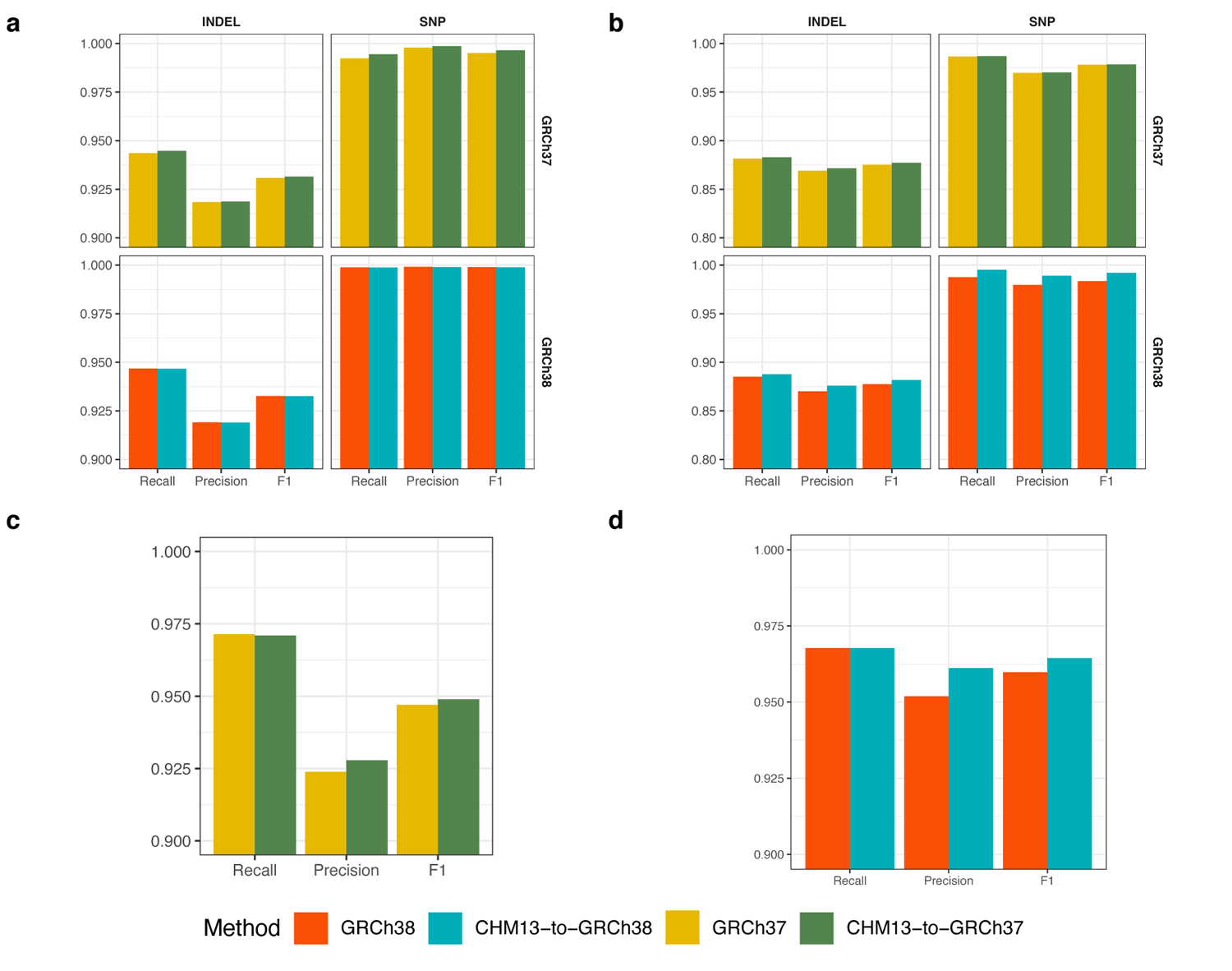
Small and structural variant calling using PacBio-HiFi reads from HG002. **a, b**, Small variant calling using minimap2–DeepVariant in **a** GIAB v4.2.1 regions and **b** GIAB CMRG regions. **c**, Structural variant calling in GRCh37 GIAB Tier 1 benchmark regions. **d**, Structural variant calling in GRCh38 CMRG regions.

Using the same PacBio-HiFi dataset, we called structural variants (SVs) using Sniffles2^36^ and analyzed the results using truvari^37^ (Section 4.5.2). We evaluated the SV calls using the GIAB Tier 1 benchmark regions for GRCh37^28^ and the GIAB CMRG benchmark for GRCh38^26^. Note that there is not a genome-wide SV benchmark for each of the references. CHM13-to-GRCh37 removed 44 (5.7%) false-positive SV errors (FPs) compared to direct-to-GRCh37 (Figure 4c and Table S11). Most FPs were shared between both methods, and the majority of these were insertions enriched in low-complexity regions. Many of the resolved FPs associated with regions having a many-to-1 mapping from donor to target, or “mapping collapse” (Figure 6b). While CHM13-to-GRCh37 resulted in 4 more false-negative SV errors (FNs) compared to direct-to-GRCh37 using the GIAB Tier 1 SV calls, we observed examples where the GIAB calls did not agree with haplotype-resolved assemblies for the same individual^38^ (Figures S6 and S7). We visually inspected these examples and observed evidence of mapping collapse including abnormal coverage and loci with more than two haplotypes, suggesting improved mapping and reduced SV FNs by levioSAM2. Compared to direct-to-GRCh38 in the GIAB CMRG regions, CHM13-to-GRCh38 removed 2 (11.8%) SV calling errors. Both were false deletions in the *KMT2C* gene (Figure 4d and Table S12).

### LevioSAM2 reduces large-scale mapping artifacts

LevioSAM2 can resolve or reduce large-scale mapping artifacts compared to direct-to-GRC pipelines. We first examined the small variant calls using the real 30× HG002 short read dataset. In the medically relevant gene *KMT2C*, we observed high-density mapping errors which had been reported by the GIAB CMRG study^26^. The errors were due to 15-kbp sequences in *KMT2C* in HG002 but collapsed in GRCh37. Therefore, sequences from both regions mapped to *KMT2C*, resulting in an abnormally high mapping depth (up to 296×) and a high alternate allele density within *KMT2C*. The missing homolog was assembled in T2T-CHM13 and marked as a suppressed region by the levioSAM2 annotation workflow. Mapping to T2T-CHM13 correctly placed the reads in the appropriate locus and the levioSAM2 workflow resolved most mapping errors in this region (Figure 5a). We also observed that DeepVariant reported a low number of variants in this region using GRCh37, even when the variant allele frequencies were as high as 0.8. We reasoned that DeepVariant learned this signature of mapping artifacts and suppressed variants in its model (Figure S8 and Note S1).

**Figure 5:**
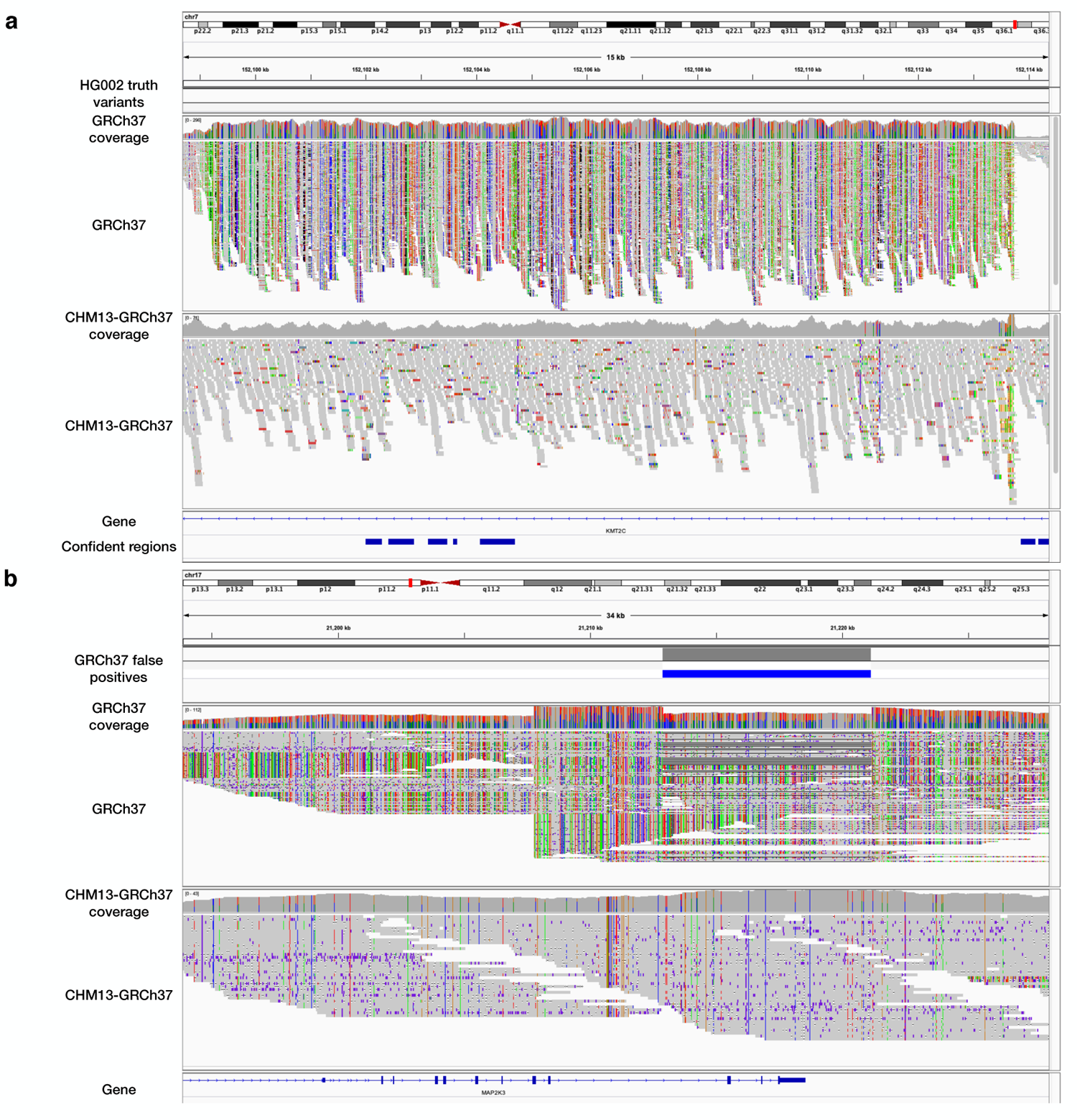
LevioSAM2 resolved large-scale mapping errors in medically relevant genes. **a**, Mapping short reads to gene *KMT2C*. **b**, Mapping PacBio HiFi long reads to gene *MAP2K3*. The “GRCh37” track in both plots **a** and **b** show high mapping depth and high density of alternate alleles (bars with non-gray colors), suggesting collapsed mapping.

We then analyzed mappings of the HG002 PacBio-HiFi long read dataset and observed similar large-scale mapping improvements by levioSAM2. Similar to the short-read example in gene *KMT2C*, we observed a high density of alternate alleles in a 56-kbp region which overlapped with another medically relevant gene, *MAP2K3*^39^. We showed that CHM13-to-GRCh37 significantly reduced mapping depth and alternate allele density in this region, suggesting improved mapping. When assessed with the GIAB Tier 1 benchmark for SVs, CHM13-to-GRCh37 avoided a false structural deletion (8.2 kbp) (Figure 5b).

### LevioSAM2 is computationally efficient

LevioSAM2-lift’s bitvector-based algorithm is fast and memory-efficient (Figure 6a). Compared to CrossMap^15^, levioSAM2-lift used 20.1% of the wall-clock time and 19.9% the peak memory (64.3 MB vs. 341.9 MB) when run on a single thread. Unlike CrossMap, levioSAM2-lift supports multi-thread processing and used only 7.3% wall time and 19.4% of the memory (66.4 MB vs. 341.9 MB) when using 4 threads (see Figure S4 for thread scaling). LevioSAM2-lift was able to replicate the results of CrossMap, but includes several additional features that are important in practice. For instance, levioSAM2 can lift mappings spanning chain-file gaps and can update CIGAR-string information in the output mappings, enabling lifting of long-read mappings. Further, levioSAM2-lift can optionally update edit distance (`NM:i`) and mismatch encoding alignment string (`MD:z`).

**Figure 6:**
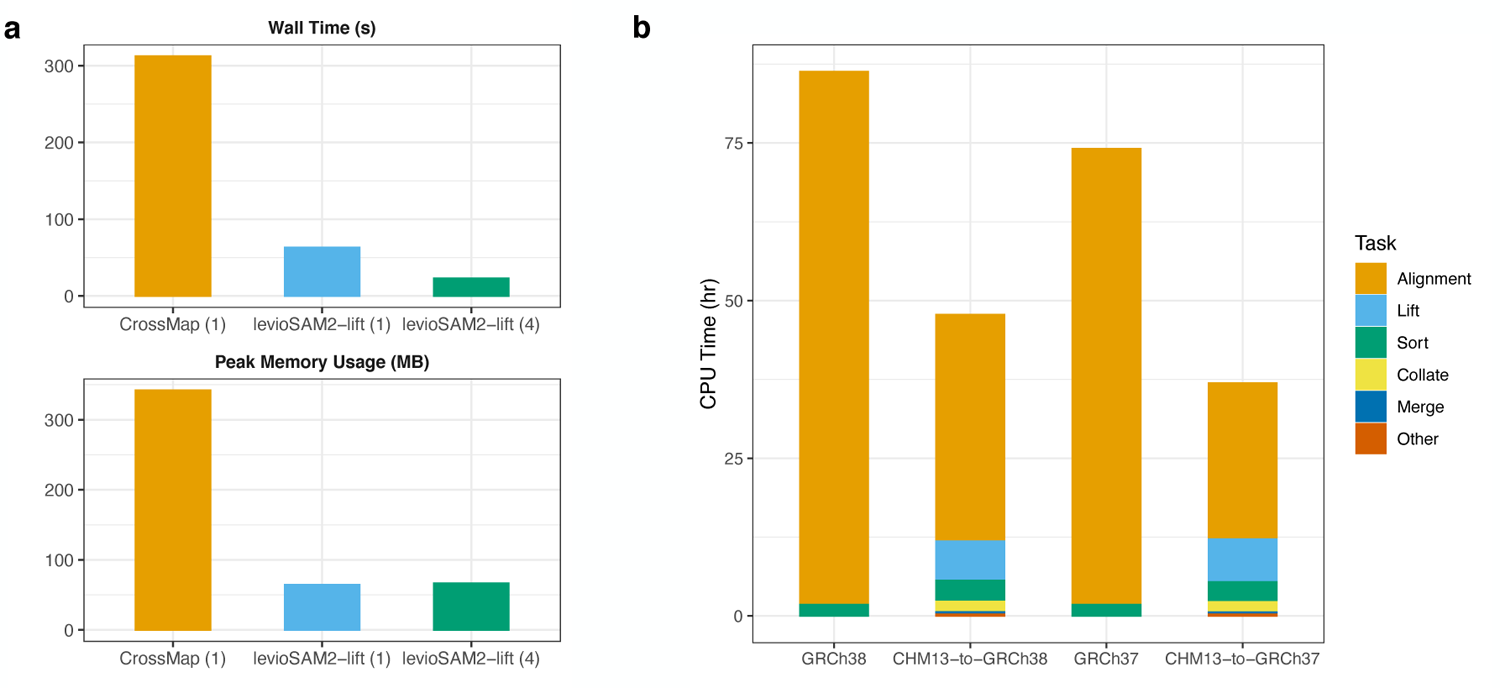
LevioSAM2 is computationally efficient and accurate when lifting from T2T-CHM13 compared to a direct mapping pipeline. **a**, Computational efficiency of methods that support lift-over of alignments for a 0.3× paired-end WGS dataset. Numbers in parentheses show the number of threads used. **b**, CPU time usage of mapping using BWA-MEM vs. lifting over using levioSAM2 for a 30× paired-end WGS dataset. In the lift-over tasks, mapping to T2T-CHM13 was not included in the runtime measurement.

The full levioSAM2 workflow (excluding the initial mapping step to T2T-CHM13) was also faster than a standard approach that simply maps all the 30 Illumina pairedend WGS reads for HG002 to the target reference (Figure 6b). We compared the CPU time used by BWA-MEM^31^ when aligning directly to the target and by the samtools sort command^40, 41^ to the corresponding levioSAM2 run. For a 30× real WGS dataset initially mapped to T2T-CHM13, the levioSAM2 workflows took 36.9 CPU hours for CHM13-to-GRCh37 and 47.6 CPU hours for CHM13-to-GRCh38. Compared to directly mapping to the target genome, the CHM13-to-GRCh37 method took 49.8% of the time (74.1 hours) and the CHM13-to-GRCh38 method took 55.4% of the time (86.3 hours). While other stages in the levioSAM2 workflow used more memory compared to levioSAM2-lift, they didn’t use more than 35 GB memory (Note S2 and Figure S3).

## 3 Discussion

LevioSAM2 lifts mappings from a source reference to a target reference while selectively remapping the subset of reads for which lifting is not appropriate. LevioSAM2 uses efficient succinct data structures and is much more efficient than existing tools. While it is common to lift post-read-mapping results like variant calls between references^14, 15^, these translations can be inaccurate due to large-scale differences between references^12, 18–20^. Starting from a BAM file mapped to T2T-CHM13, levioSAM2 generated GRC-based mappings 44.7-50.2% faster than a method which remapped all the reads. Mapping and genotyping results after using levioSAM2 were more accurate than those obtained using a single reference in both simulated and real-data scenarios. Since levioSAM2 uses linear references, its results are readily interpretable using common tools like the Integrative Genomics Viewer (IGV)^42^.

The small-variant calling improvements by levioSAM2 were enriched in difficult regions with presence of low-mappability units, segmental duplications, and assembly artifacts. These regions include the challenging medically relevant genes (CMRG), where accurate variant calling is known to be difficult. LevioSAM2 also improved long read mapping, demonstrated by more accurate small- and structural-variant calling. Notably, levioSAM2 resolved most mapping errors in a 15-kbp region in the *KMT2C* gene and a 56-kbp region in the *MAP2K3* gene in GRCh37.

As the need to move alignments between assemblies becomes more common, it will be important to improve whole-genome alignment maps between those references. While the UCSC “lift over construction” recipe^14, 43^ has become a standard approach, more work is needed to assess the quality of the resulting chain files^44^. For example, sequences with multiple copies are challenging to accurately place, and some chain files do not guarantee 1-to-1 mapping between the source and target references. While the levioSAM2 framework tolerates errors in a chain-file alignment, we expect improved whole-genome maps to further enhance the accuracy and computational performance of levioSAM2.

LevioSAM2’s decisions on whether to commit, defer or suppress a read mapping rely on relatively simple heuristics. We have shown these are effective in reducing the overall number of needed re-alignments. Nevertheless, we expect that levioSAM2’s accuracy can be further improved by making these decisions more data- and model-driven, perhaps using properties of the input such as the read length, assay type, or data quality.

The future may see a shift toward more sophisticated pangenome representations, e.g. made up of high-quality population-scale genome assemblies^35, 45^. We expect that accurate and efficient lift-over methods like levioSAM2 will be useful in these contexts as well. While construction of a pangenome graph can require use of expensive multiple- or progressive-alignment algorithms, the levioSAM2 approach can be applied to any pair of genomes with a whole-genome alignment to each other (in the form of a chain file).

Rapid progress in genome assembly and sequencing is making hundreds of high-quality reference assemblies available^35^. These assemblies provide demonstrated improvements for short and long-read variant calling^2, 4^. However, a rich set of annotations, key for interpretation, are built on previous references and difficult or impossible to translate to every new assembly. LevioSAM2 enables analyses that have the best of both worlds by improving mapping for the original reference without losing all its secondary information while also providing mappings for novel genomic discovery in regions unique to the new reference.

## 4 Methods

### 4.1 Efficient lift-over using succinct data structures

A chain file describes a pairwise whole-genome alignment^14^. It consists of many “chains”, each a set of co-linear alignments between the source and the target reference. An alignment is further split into an interleaved sequence of aligned segments and gaps. An aligned segment can include matches and mismatches but not gaps. A gap can appear in one of the references or both. Each line in a chain specifies one aligned segment and up to two gaps (Figure 1c: bottom).

LevioSAM2 first sorts the aligned segments by position and stores them in a chain interval array. Each chain interval records information useful for lifting, including target contig, strand, and offset. The offset is represented as the difference between source position and target position.

To enable queries against the chain interval array, levioSAM2 builds a pair genome-length of succinct bit vectors (“BV”) (Fig. 1c: top). It uses the `start_bv` bit vector to encode starting positions of all chain intervals and uses the `end_bv` vector for ending positions. Both bit vectors are supplemented with data structures enabling constant-time rank queries, as provided by the SDSL library^46^. The levioSAM2 implementation further wraps these bit vectors and arrays in an unordered map, with source contigs as keys to the map.

When querying a position, levioSAM2 first locates the index of the corresponding chain interval by performing a `rank` query over both `start_bv` and `end_bv`. A rank query computes the number of set bits (bits equal to 1) prior to the queried position. If a position *p* is within a chain interval, it must be that `rank`_start_bv_(*p*) `rank`_end_bv_(*p*) = 1. For a position outside of any chain interval, levioSAM2 checks the distance between the position and its neighbor chain interval boundary. If the distance is under a user-defined threshold, the query is assigned to the neighbor. LevioSAM2 queries the chain interval array using the index and updates the contig, strand and position information. Since it is dominated by the rank query, this is a constant-time algorithm (*O*(1)), which is faster than the commonly-used interval tree-based algorithms^14, 15^, which can use *O*(log(*m*)) time *m* is the number of intervals.

It is necessary to update the CIGAR information of an alignment when it overlaps a chain gap. This kind of CIGAR update is not performed by CrossMap^15^. While levioSAM does perform such an update^17^, this takes *O*(*n*) time where *n* is the length of a read. Here we describe a new algorithm (Alg. 1) to update the CIGAR string. The algorithm maintains a list of chain gaps, and updates the chain gaps into the CIGAR string. Since the algorithm only traverses through the CIGAR string and gaps collection once, its time complexity is *O*(*r* + *g*), where *r* is the number of runs in the original CIGAR and *g* is the number of overlapped chain gaps in the alignment. Usually *r* and *g* are much smaller than the read length *n*, so the proposed algorithm is substantially faster than levioSAM’s. While levioSAM2 can lift over mappings spanning gaps in a chain file and update the CIGAR strings accordingly, the resulting mapping does not always align optimally after lift-over. That is, a slightly different decision about how to arrange gaps and mismatches might be more optimal. To achieve optimal alignment, levioSAM2 includes an optional realignment module that uses a localized dynamic-programming algorithm to refine the alignment. Note that we use the term “mapping” for the task of determining where the read camp from, and the term “alignment” for the more detailed task of lining individual read bases up with reference bases. We use the ksw2 library^47^ for efficient dynamic-programming-based realignments. We support parameter presets in the YAML format^48^ for popular aligners including Bowtie 2, BWA MEM, and minimap2. LevioSAM2 uses the data structures provided by htslib^49^ to process SAM and BAM files.

### 4.2 LevioSAM2 workflow with selective re-mapping

To improve mappings in the presence of large-scale differences between source and target references, levioSAM2 uses a selective re-mapping strategy. Reads that can be lifted with high confidence are “committed” and the lifted alignment is taken as the final alignment. Reads belonging to a region that is specific to the source reference are “suppressed” and left unaligned with respect to the target reference. Reads for which the lift is lower-confidence are “deferred” and are re-mapped to the target reference (Fig. 1b; Section 4.2.1). For paired-end reads, we use the “levioSAM2-collate” step to ensure deferred reads are re-mapped as a pair (Section 4.2.2). After re-mapping, “levioSAM2-reconcile” compares the re-mapped and lifted mappings for each deferred read and selects the one with higher confidence (Section 4.2.3). Committed and reconciled mappings are combined in the final output.

#### 4.2.1 Selective strategy

To determine if a given read should be committed, levioSAM2 examines a combination of the read’s alignment features and the reference genome’s “liftability” annotation. Alignment features include mapping status (mapped or not), mapping quality, fraction of clipped bases, edit distance (the `NM:i` tag), alignment score (the `AS:i` tag), and fragment length (if part of a pair). For BWA-MEM (local alignment), we used a MAPQ cutoff of 30 or an alignment score cutoff of 100; for Bowtie 2 (end-to-end alignment), we used a MAPQ cutoff of 10 or an alignment score cutoff of −10.

**Algorithm 1:**
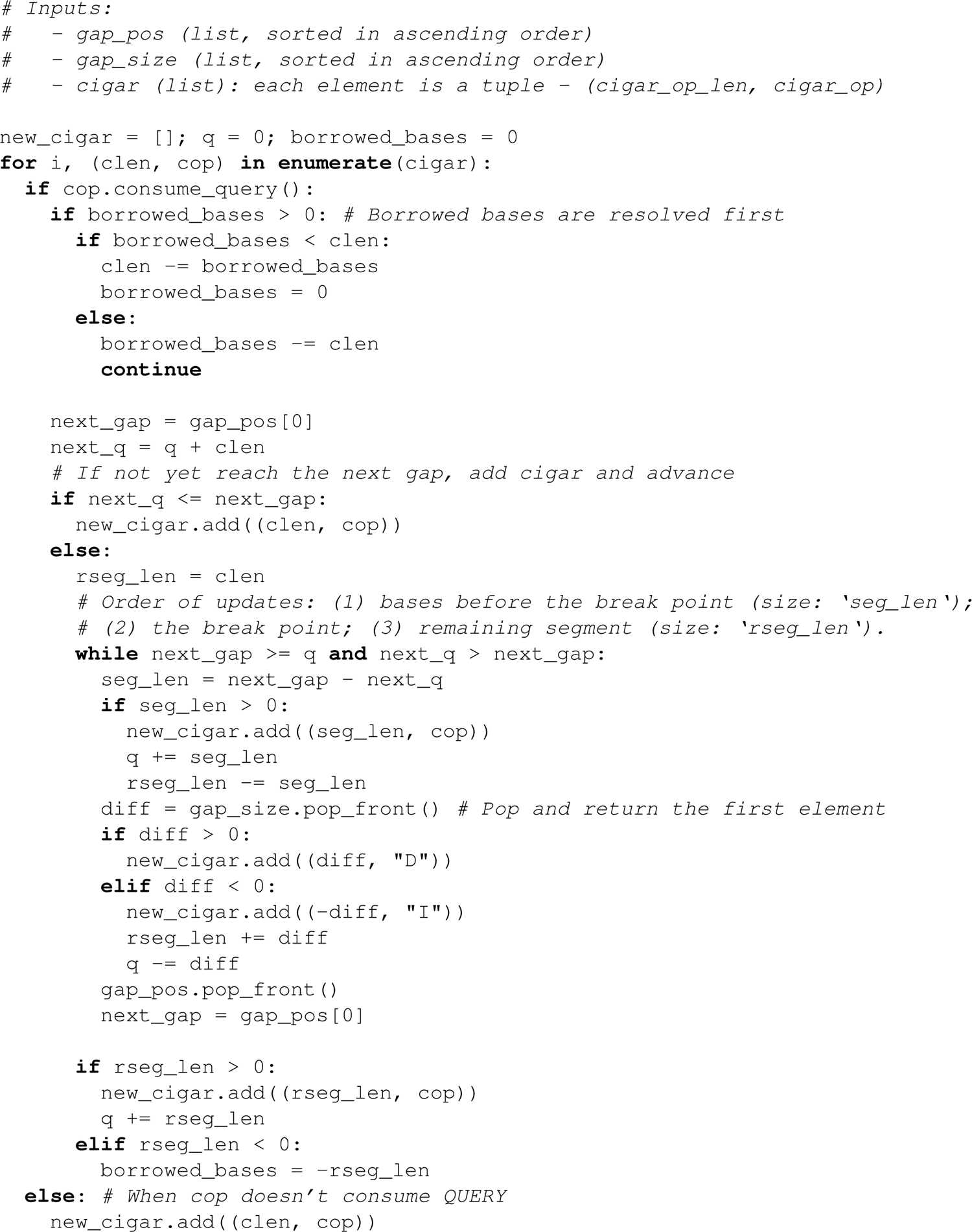
UpdateCigar

The genomic liftability annotation is a BED file marking some regions as “unliftable” and some as “mappability-reduced.” The annotation is generated ahead of the lift-over process and is indexed using interval trees for fast queries. The “unliftable” annotation is given to regions that are unique to the source reference, like T2T-CHM13’s centromeric regions, which have no counterpart in GRCh38. Reads mapping to these regions are suppressed in order to avoid false re-mappings. Suppression tends to improve computational performance as well, since suppressed reads often come from repetitive regions and require a disproportionate amount of alignment effort. To determine which regions are unliftable, we first extract sequences from the source reference that are not in the chain file and are longer than 5000 bps. We then align the sequences to the target reference using Winnowmap2^34^ and label either unmapped or repetitive (mapping to multiple loci in target) regions as unliftable.

LevioSAM2 also looks for “mappability-reduced” regions in the source to gauge confidence in lifted mappings. Mappability-reduced regions have high mappability in the source reference but lower mappability after being mapped to the target. To build the mappability-reduced annotations, we use GenMap2^50^ to calculate mappability for both source and target references (using 100-mers and a 0.01 mismatch-rate tolerance). We extract uniquely mappable regions in the source reference and lift them to the target. We then overlap these lifted unique regions with the low-mappability regions in the target. Reads lifted to these regions are deferred and re-mapped.

#### 4.2.2 Collate

For paired-end reads, the selective strategy can result in cases where the paired ends are assigned to different groups. Re-mapping accuracy for single-end reads is usually lower than for paired-end reads, especially when the reads overlap indels. Thus, we develop the “levioSAM2-collate” method to additionally defer the mate of the deferred singletons, generating properly paired deferred read sets.

LevioSAM2-collate starts by reading the deferred alignments and storing the first and the second segments separately in a pair of hash maps^51^. For a properly-paired read (i.e. with both ends in the deferred group), we write both ends to a paired-deferred BAM file and remove them from the hash tables. Once all deferred alignments have been processed, any paired ends remaining in the hash maps are singletons. We then read the committed alignments and extract the reads which pair with the deferred singletons.

#### 4.2.3 Reconciling

Motivated by the “reference flow” mapping strategy^22^, we designed a “levioSAM2-reconcile” method to improve accuracy. LevioSAM2-reconcile compares the lifted and re-mapped deferred mappings, selecting the one which has higher confidence. The process starts with sorting both deferred BAM files by query name. We select the mapping with lower edit distance with respect to the target reference. Ties are broken by taking the mapping with higher mapping quality, or by making a random choice if they are still tied. By reconciling and selecting the alignment in this way, levioSAM2 can better leverage the genetic diversity provided by the additional reference.

### 4.3 Generating chain files

We used nf-LO^43^ with the minimap2 mode to build the chain files for both CHM13-to-GRCh37 and CHM13-to-GRCh38:

`nextflow run main.nf --source source.fa --target target.fa \`

`--outdir out_dir -profile local --aligner minimap2`

### 4.4 Evaluation using simulated sequencing datasets

We used the GRCh38-based SNPs for NA12878/HG001 from the 1000 Genomes Project (1KGP)^12^ to build personalized chromosome 20 and 21 references for simulation. We masked the regions that could not lift to T2T-CHM13 using bedtools-maskfasta^52^. We used the mason simulator^29^ to simulate 10M 100-bp paired-end reads (5M pairs) for both chromosomes:

`mason_simulator --num-threads 16 -ir ref.fa -n 5000000 \`

`-o out-R1.fq -or out-R2.fq -oa out.sam -iv sample.vcf`

We mapped the reads using both unmasked GRCh38 and CHM13-to-GRCh38 using default parameters for both Bowtie 2^30^ and BWA-MEM^31^. We compared mappings from the direct-to-GRCh38 and CHM13-to-GRCh38 strategies by measuring the fraction of correct mappings, where a mapping was considered correct if its leftmost mapped position (the first un-clipped base) was within 10-bp of that of its simulated origin.

### 4.5 Evaluation using real sequencing datasets

#### 4.5.1 Small variant calling and evaluation

We evaluated three real 30× coverage datasets sequenced using Illumina Novaseq and a PCR-free protocol. We also evaluated a dataset sequenced with PacBio-HiFi with 28× coverage (Table 1). The samples were from distinct ancestries, including Utah/European (HG001), Ashkenazi Jewish (HG002), and Han Chinese (HG005).

For the Illumina datasets, we mapped the reads to GRCh37, GRCh38, and T2T-CHM13 using BWA-MEM^31^ with default parameters and sorted by genomic positions using samtools^41^. We used the levioSAM2 workflow to lift the reads mapped to T2T-CHM13 to GRC references and generated CHM13-to-GRCh37 and CHM13-to-GRCh38 mappings. We then called small variants using GATK HaplotypeCaller^23^ and DeepVariant (the “WGS” model)^24^ for GRCh37, GRCh38, CHM13-to-GRCh37, and CHM13-to-GRCh38. For the PacBio-HiFi dataset, we followed similar procedures, except we used minimap2^33^ as the read mapper and called small variants using the haplotype-sorting DeepVariant pipeline with the “PACBIO” model^53, 54^.

For both sequencing data types, we evaluated the accuracy of small variants using hap.py^55^ and the Genome in a Bottle (GIAB) v4.2.1^25^ and the GIAB Challenging Medically Relevant Gene (CMRG) benchmark^26^ truth sets. Hap.py reports small variant calling accuracy measures stratified by variant type. We sometimes reported the overall (SNP and indel) *F*_1_ of one dataset for simplicity. We did so by adding the true positives (TP), false positives (FP), and false negatives (FN) for SNPs and indels, then calculating overall *F*_1_ with 2 TP*/*(TP + 0.5(FP + FN)). Similarly, for the *F*_1_ of multiple datasets, we summed the measures for all datasets and calculated the *F*_1_.

#### 4.5.2 Structural variant calling and evaluation

We called structural variants (SV) for the PacBio-HiFi HG002 dataset using Sniffles2 v2.0.1^36^. We used the “germline SV calling” mode with default parameters, without providing any tandem repeat annotations. We next compressed the VCF files for each dataset using bgzip and indexed them with tabix^49^. Finally, we benchmarked and compared the SV calls using the GIAB Tier 1 benchmark regions for GRCh37^28^ and the GIAB CMRG benchmark for GRCh38^26^ using truvari 2.1^37^ and following the GIAB benchmarking instructions.

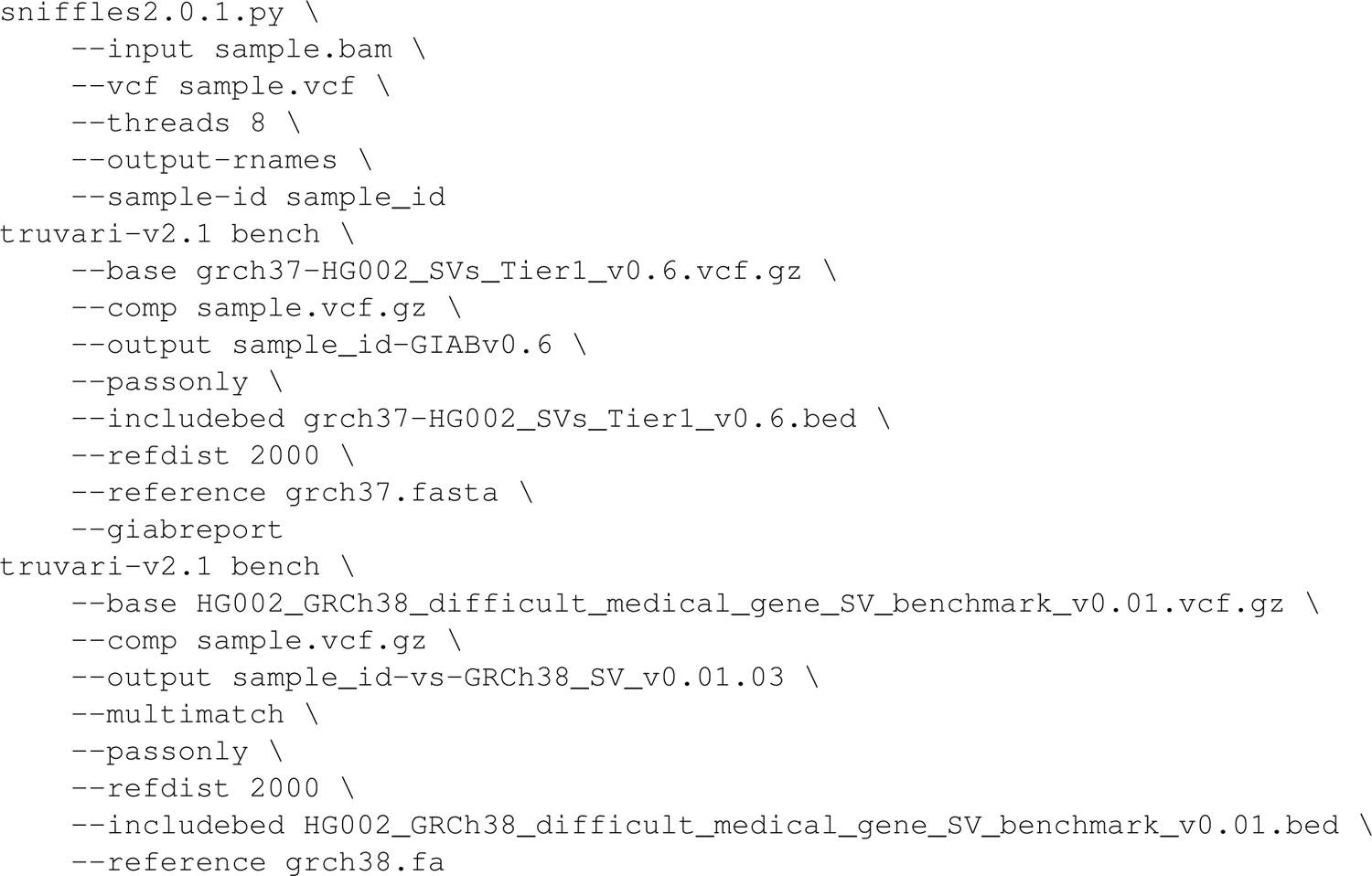

### 4.6 Computational efficiency measurement

The computing nodes we used were Intel Cascade Lake 6248R with 48 cores per node and 192 GB of memory. We requested 36 cores for all experiments except for the thread scaling experiments. We used GNU Time^56^ to measure CPU time (“User time” + “System time”), wall clock time (“Elapsed (wall clock) time”) and peak memory usage (“Maximum resident set size”).

## 5 Availability of data and materials

The software is available at https://github.com/milkschen/leviosam2 under the MIT license. The experiments described in this paper are described at https://github.com/milkschen/levioSAM2-experiments under the MIT license.

## Acknowledgements

Part of this research project was conducted using computational resources at the Maryland Advanced Research Computing Center (MARCC). Pre-built levioSAM2 resources for T2T-CHM13 to GRC references are made freely available on Amazon Web Services thanks to the AWS Public Dataset Program. We thank Taher Mun for his advice and contribution to the levioSAM2 programming infrastructure. We appreciate advice from Hann-Chyun Chen on software deployment, Christopher Pockrandt on mappability resources, Alaina Shumate on gene lift-over, and Samantha Zarate on T2T-CHM13 variant analysis. We also thank Andrew Carroll and Pi-Chuan Chang for DeepVariant discussions, Arang Rhie for T2T-CHM13 discussions, and Justin Zook for GIAB strata suggestions.

## Authors’ contributions

NC, SK, AMP, and BL designed the method. NC wrote the software. NC and LFP performed the experiment. NC, LFP, FS, SK, AMP, and BL performed analysis and wrote the manuscript. All authors read and approved the final manuscript.

## Ethics approval

Not applicable.

## Consent for publication

Not applicable.

## Supplementary Notes

### S1 DeepVariant calls in difficult-to-map regions

We examined the DeepVariant calls in the *KMT2C* gene, where there were known mapping collapses for HG002 data when using direct-to-GRCh37 (see Results). We noticed that DeepVariant made many homozygous reference calls even when the variant allele fraction (VAF) were as high as 0.82 (Figure S8). The GIAB truth set reported few variants in this region. We reasoned that DeepVariant could “recognize” mapping artifacts and adjust its decisions in difficult-to-map regions.

### S2 Computational efficiency of levioSAM2

We measured the CPU time and peak memory usage of each step in the levioSAM2 and typical pipelines (Figure 6b and Figure S3). In the levioSAM2 workflows, lifting alignments over took 19.2% CPU time (7.1 hours) for CHM13-to-GRCh37 and 13.6% (6.5 hours) for CHM13-to-GRCh38. The majority of the CPU-time usage for the levioSAM2 workflow was in the remapping step, taking 66.2% (24.5 hours; GRCh37) and 74.6% (35.6 hours; GRCh38) of time. Mapping the deferred reads took longer and had a higher memory footprint compared to the direct-to-target mapping task, likely because of the higher incidence of repetitive alignments for deferred reads. The most memory-consuming step was levioSAM2-collate, since we used a hash map to store unpaired deferred reads. For memory-limited systems, it will be straightforward to reduce the memory bottleneck with a marginal increase in CPU time.

## Supplementary Figures

**Figure S1:**
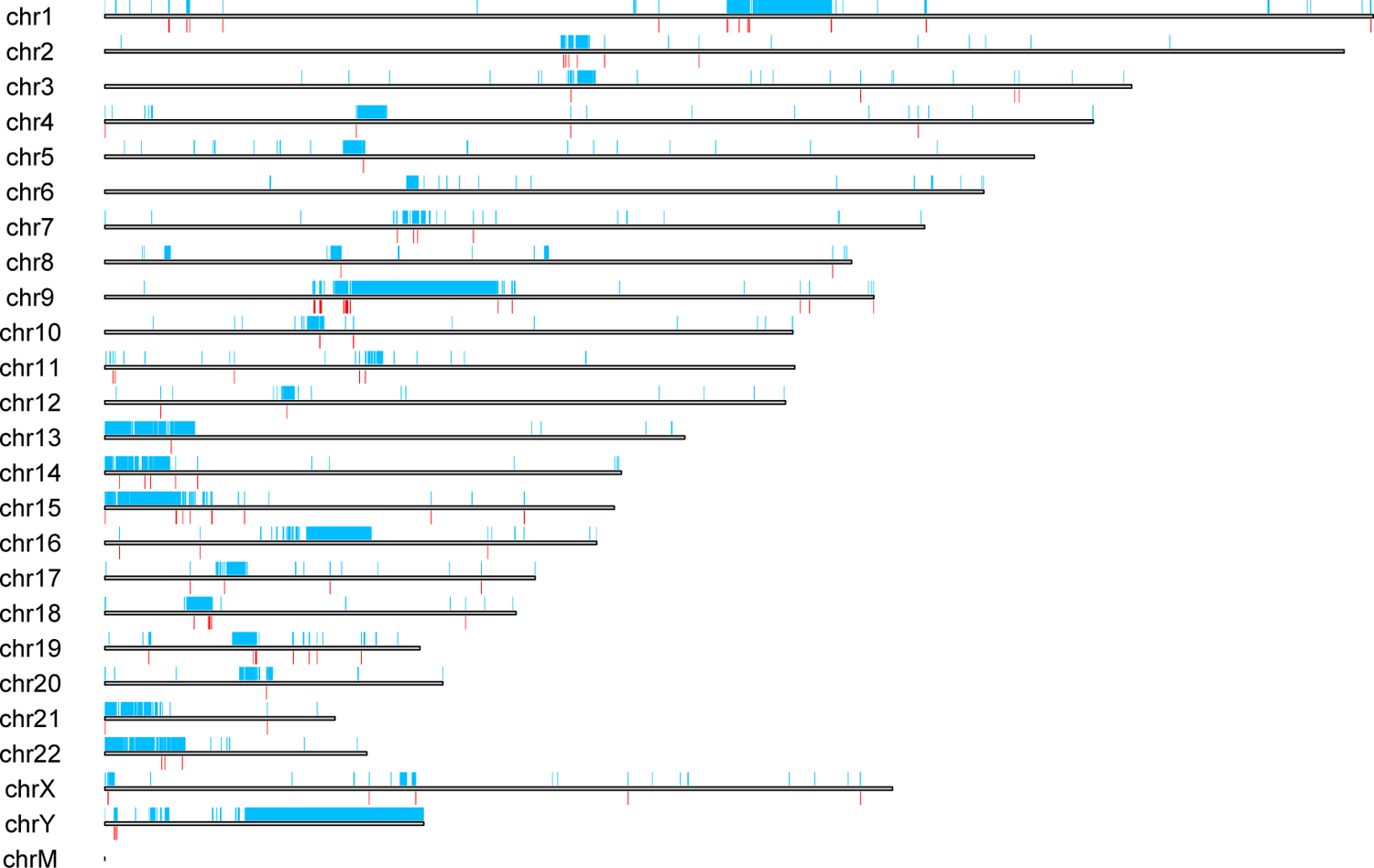
Regions unique to T2T-CHM13^1^ compared to GRCh38 (blue) and high quality calls from DeepVariant in these regions (red).

**Figure S2:**
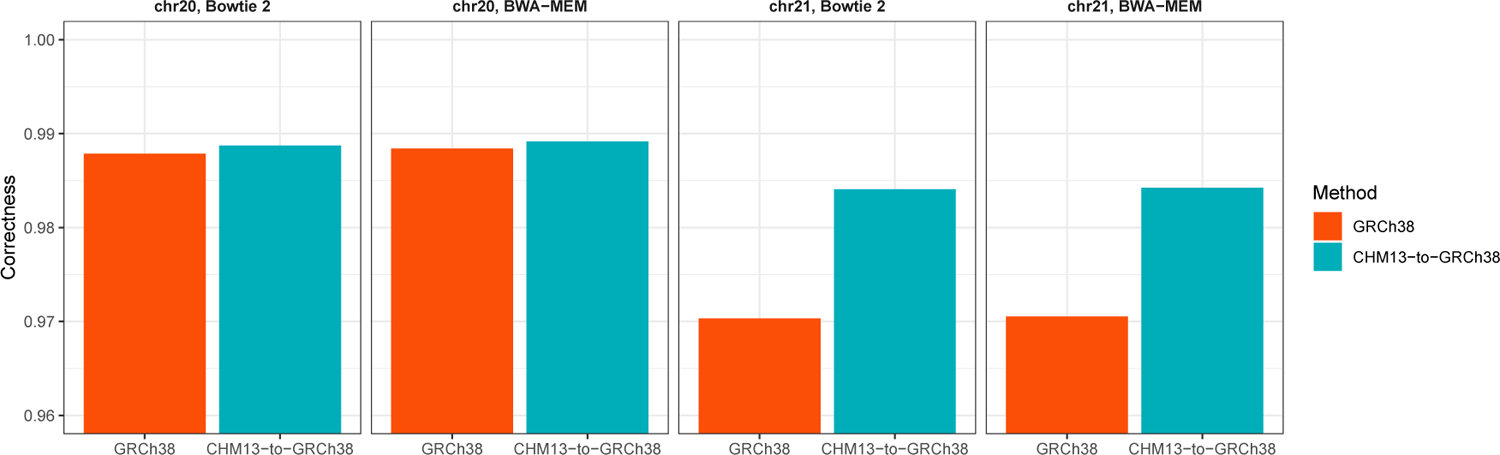
Mapping accuracy using simulated reads that carry GRCh38-based HG001 genotypes^2^.

**Figure S3:**
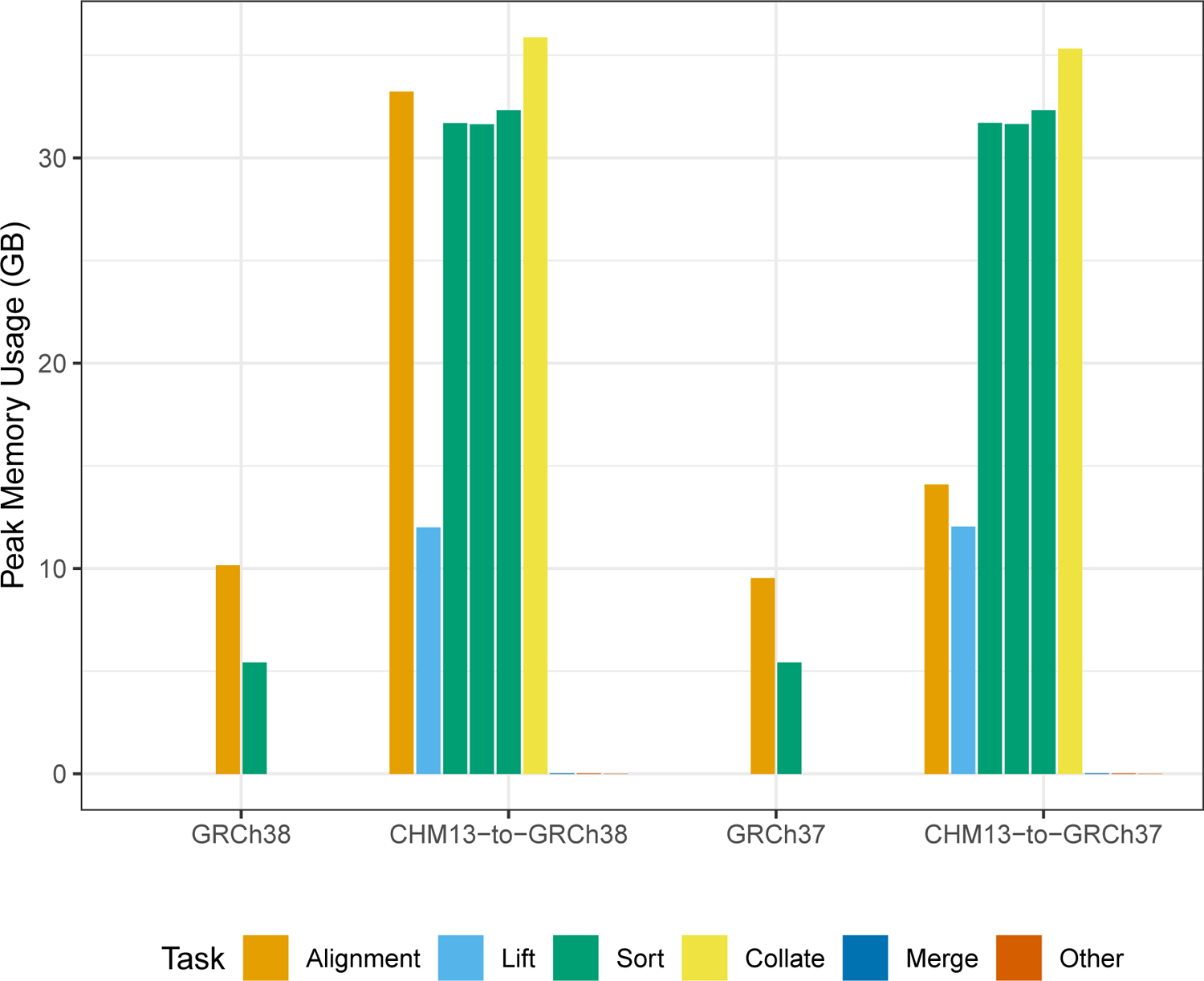
Peak memory usage of levioSAM2 and direct-to-GRC pipelines using a real 30*×* WGS dataset from HG002.

**Figure S4:**
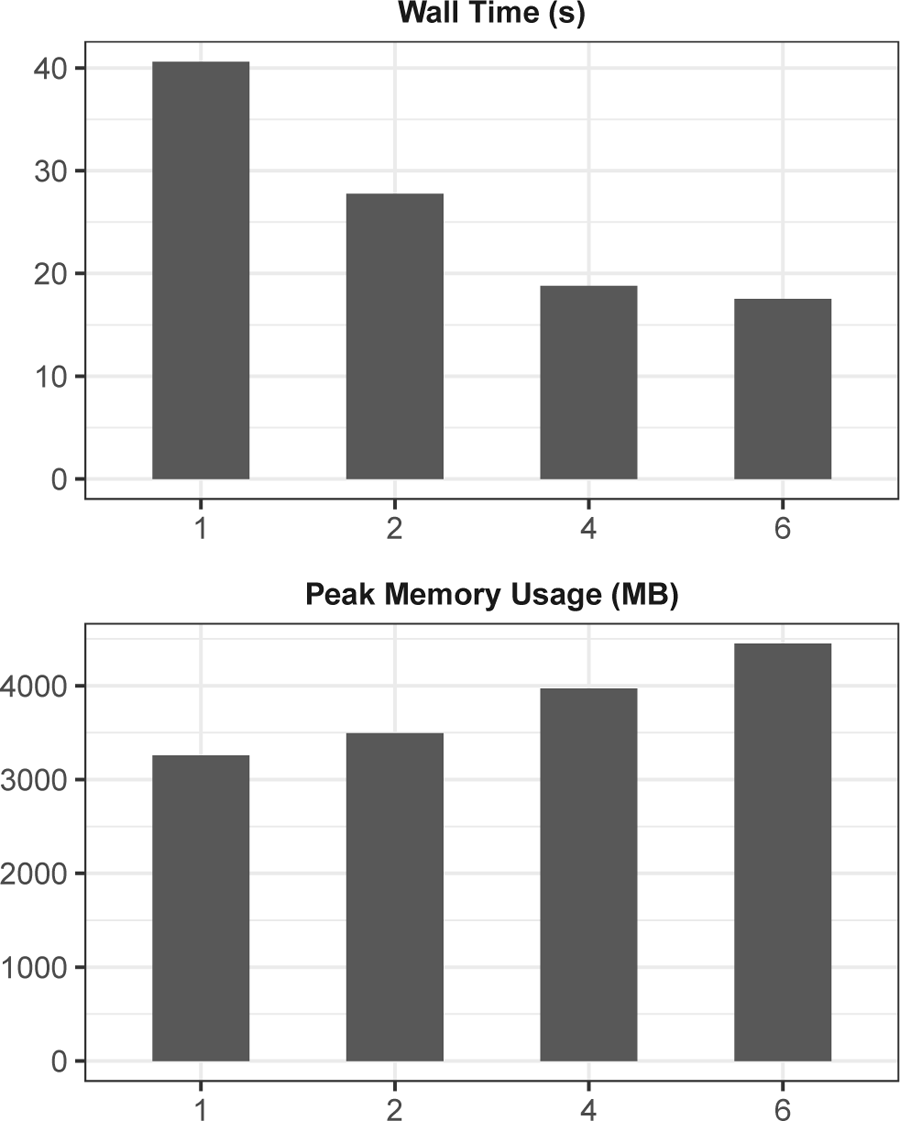
Thread scaling of levioSAM2-lift. 3.6M pairs (0.3 × coverage) of real Illumina reads from the real HG002 dataset were used. Wall clock time (second) and peak memory usage (MB) were measured using GNU Time.

**Figure S5:**
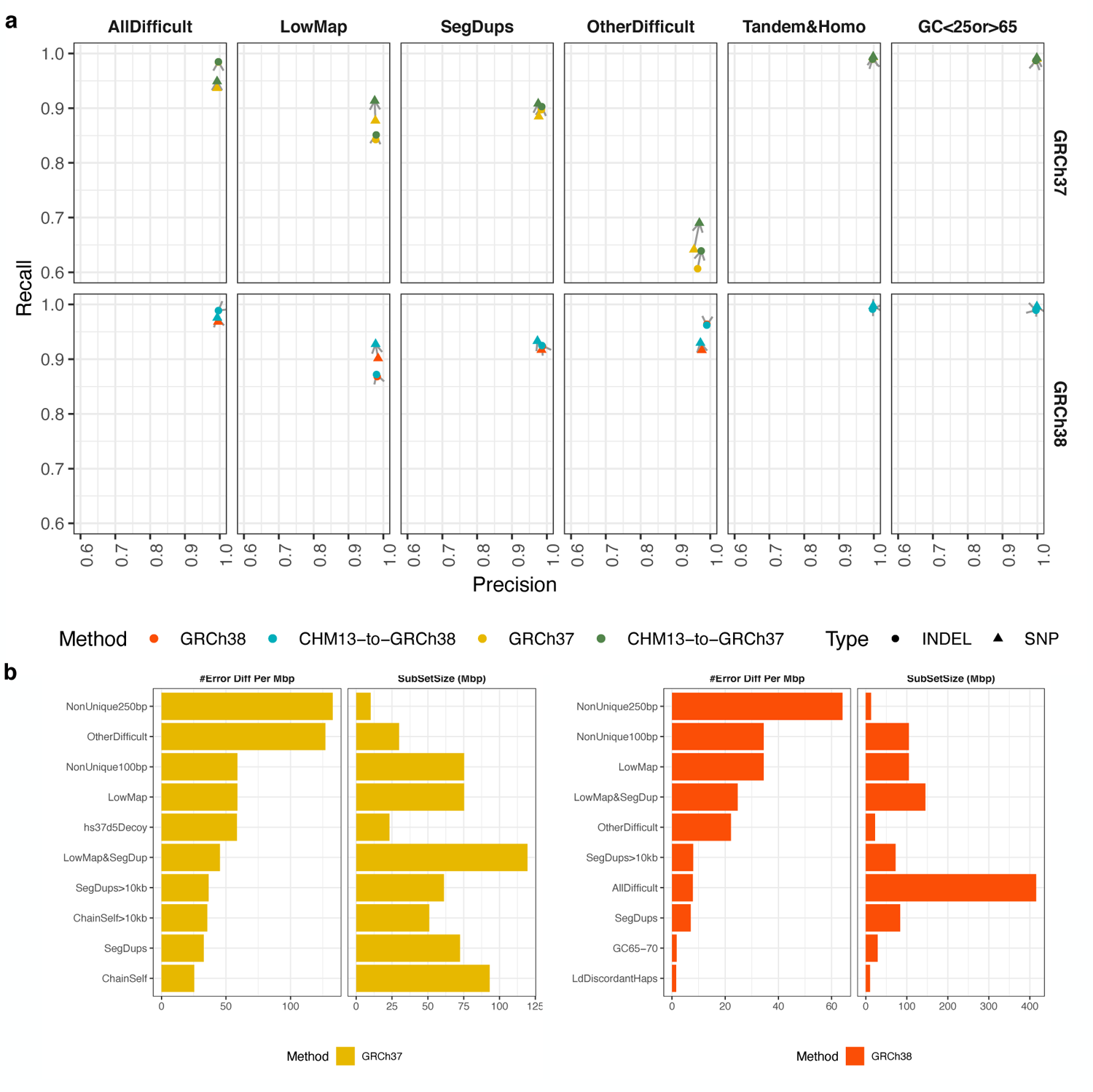
Small variant calling performance in difficult regions. **a**, Small variant calling accuracy in major difficult genomic regions for HG002. **b**, GIAB stratified regions with top small variant calling error reduction densities by levioSAM2. Small variants in both plots were called using using DeepVariant.

**Figure S6:**
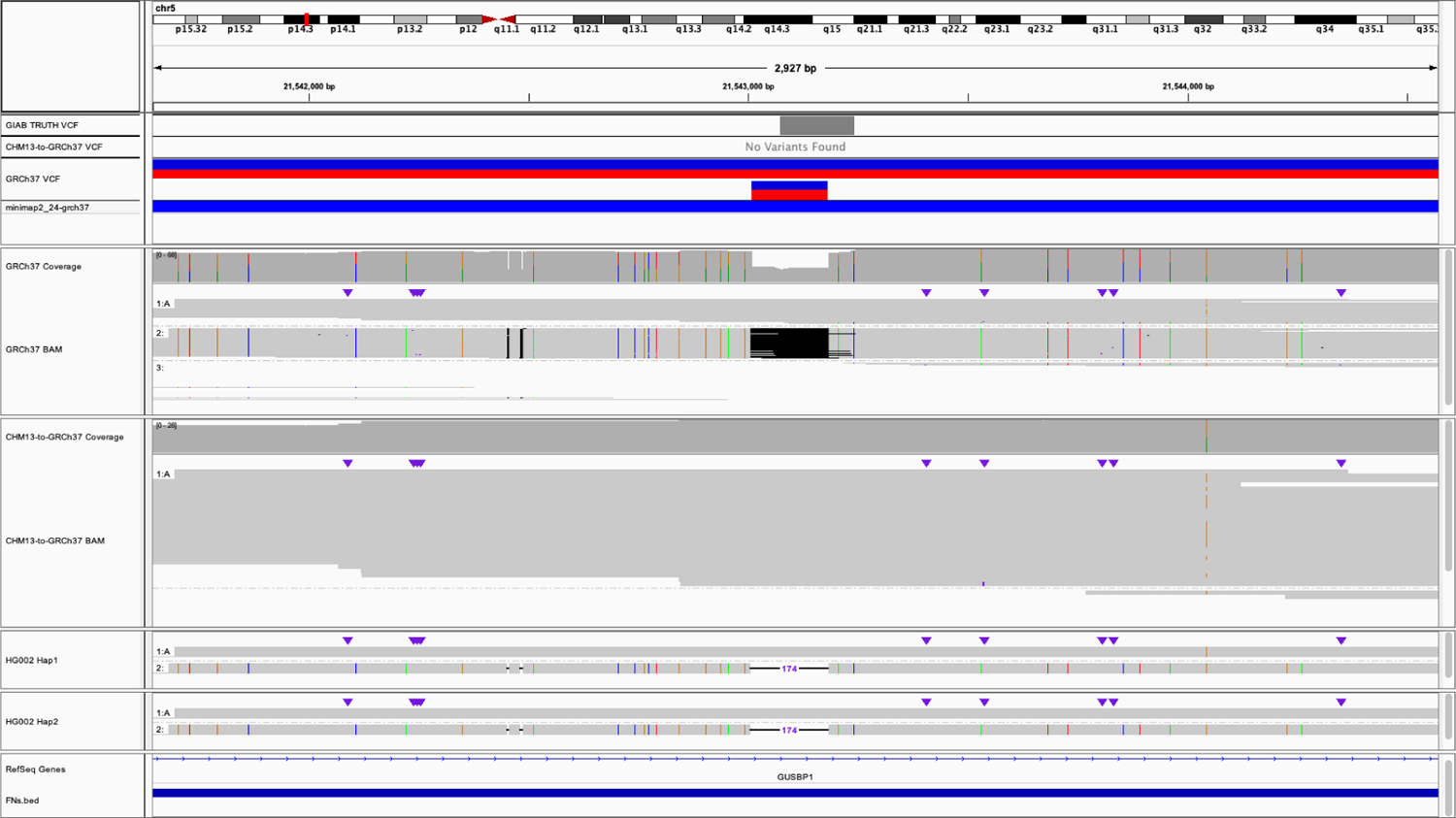
IGV visualization near chr5:21,543,010. The reads were grouped using the allele at chr5:21,543,010. A 174-bp DEL was called when using direct-to-GRCh37, matching the GIAB Tier 1 SV callset. However, personalized whole-genome assemblies suggested collapse mapping in this region and the CHM13-to-GRCh37 mappings showed better concordance with the assemblies.

**Figure S7:**
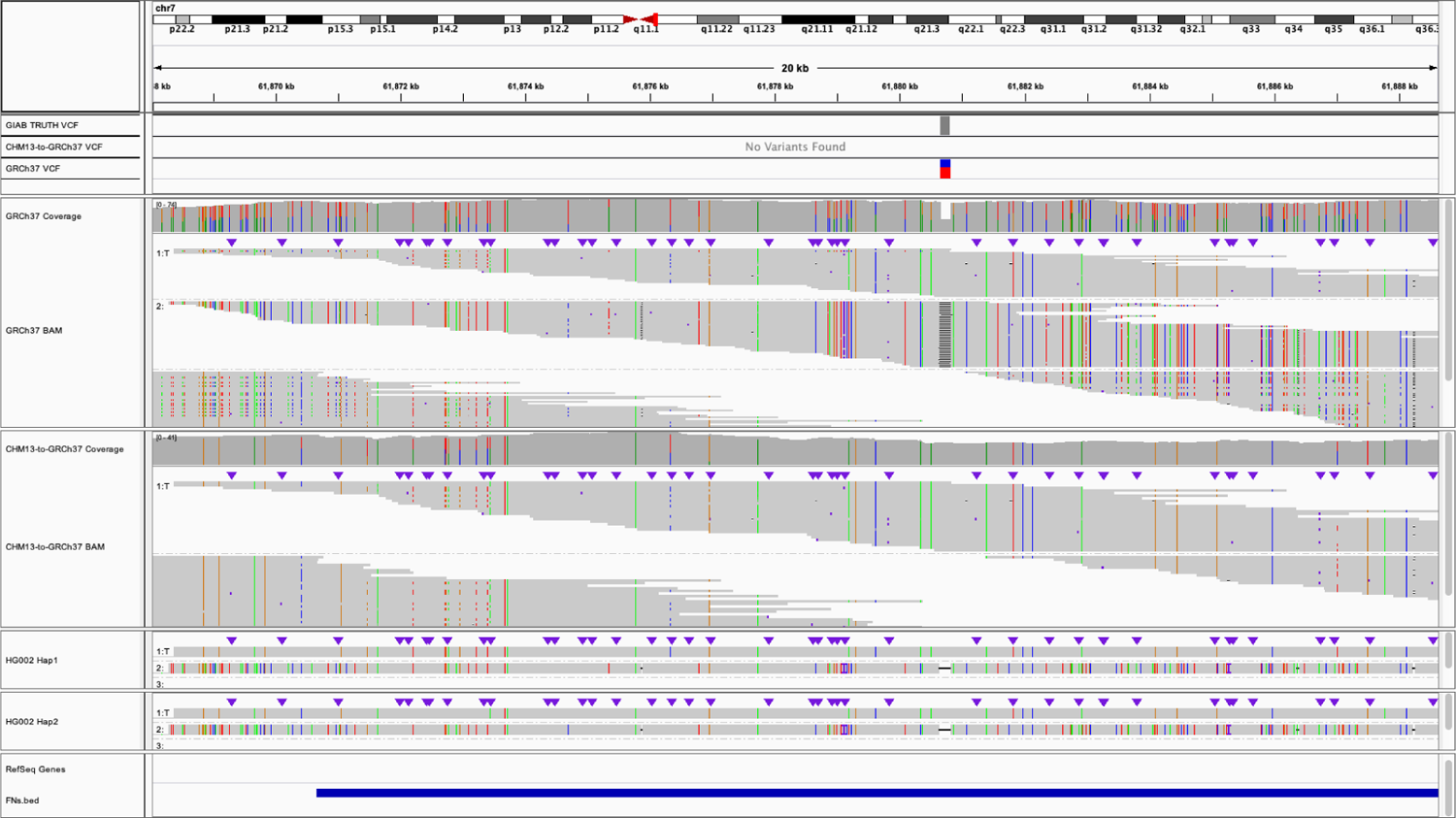
IGV visualization near chr7:61,880,665. The reads were grouped using the allele at chr7:61,880,665. A 166-bp DEL was called when using direct-to-GRCh37, matching the GIAB Tier 1 SV callset. However, personalized whole-genome assemblies suggested collapse mapping in this region and the CHM13-to-GRCh37 mappings showed better concordance with the assemblies.

**Figure S8:**
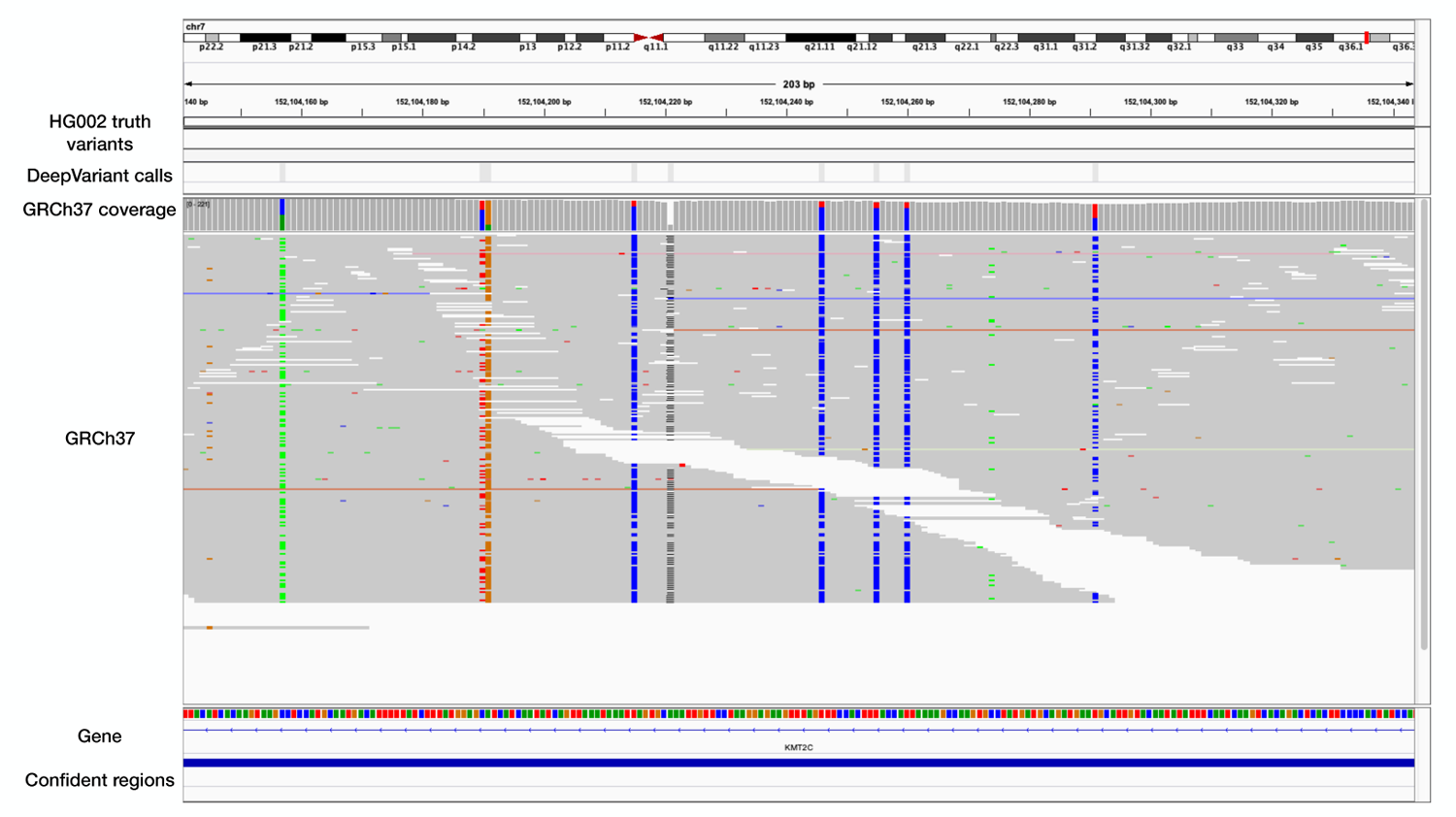
DeepVariant calls in chr7:152,104,140-152,104,343 (located in the *KMT2C* gene). This is a region annotated as high confidence (“Confident regions”) but has no truth variants (“HG002 truth variants”). Gray bars in the “DeepVariant calls” track show homozygous reference variant calls. Colors other than gray in the “GRCh37” and “GRCh37 coverage” tracks show alternate alleles.

## Supplementary Tables

**Table S1:** Software version

|  |  |
| --- | --- |
| levioSAM2 | v0.2.0 |
| BWA-MEM<sup>3</sup> | 0.7.17-r1188 |
| minimap2<sup>4</sup> | 2.24-r1122 |
| Winnowmap2<sup>5</sup> | 2.03 |
| Bowtie 2<sup>6</sup> | 2.3.5.1 |
| bedtools<sup>7</sup> | v2.30.0-48-g868a9a24 |
| GATK<sup>8</sup> | v4.2.2.0 |
| HTSJDK<sup>9</sup> | 2.24.1 |
| Picard<sup>10</sup> | 2.25.4 |
| DeepVariant<sup>11</sup> | 1.2.0 |
| Hap.py<sup>12</sup> | v0.3.8-17-gf15de4a |
| Sniffles2<sup>13</sup> | 2.0.1 |
| whatshap<sup>14</sup> | 1.2.1 |
| truvari<sup>15</sup> | v2.1 |
| nf-LO<sup>16</sup> | 1.5.1 |
| liftOver<sup>17</sup> | *accessed on Sep 14, 2021* |
| CrossMap<sup>18</sup> | v0.5.4 |
| mason2<sup>19</sup> | 2.0.9 |
| GNU Time<sup>20</sup> | 1.9 |
| IGV<sup>21</sup> | 2.6.3 |

**Table S2:** Small variant calling accuracy for 30x WGS datasets using BWA-MEM–GATK-HaplotypeCaller in all GIAB v4.2.1 regions^22^

| Sample | Method | Type | Recall | Precision | $F_1$ | TP | FN | FP |
| --- | --- | --- | --- | --- | --- | --- | --- | --- |
| HG001 | GRCh38 | SNP | 0.9932 | 0.9875 | 0.9903 | 3232144 | 22228 | 41050 |
| HG001 | CHM13-to-GRCh38 | SNP | 0.9950 | 0.9908 | 0.9929 | 3238118 | 16254 | 30230 |
| HG001 | GRCh37 | SNP | 0.9869 | 0.9790 | 0.9829 | 3209109 | 42734 | 68868 |
| HG001 | CHM13-to-GRCh37 | SNP | 0.9900 | 0.9903 | 0.9901 | 3219275 | 32568 | 31572 |
| HG001 | GRCh38 | INDEL | 0.9902 | 0.9889 | 0.9895 | 463115 | 4578 | 5418 |
| HG001 | CHM13-to-GRCh38 | INDEL | 0.9904 | 0.9907 | 0.9905 | 463195 | 4498 | 4537 |
| HG001 | GRCh37 | INDEL | 0.9863 | 0.9832 | 0.9848 | 460484 | 6391 | 8148 |
| HG001 | CHM13-to-GRCh37 | INDEL | 0.9870 | 0.9897 | 0.9884 | 460816 | 6059 | 4981 |
| HG002 | GRCh38 | SNP | 0.9922 | 0.9877 | 0.9899 | 3338776 | 26350 | 41484 |
| HG002 | CHM13-to-GRCh38 | SNP | 0.9943 | 0.9912 | 0.9927 | 3345806 | 19320 | 29656 |
| HG002 | GRCh37 | SNP | 0.9859 | 0.9802 | 0.9831 | 3305380 | 47305 | 66646 |
| HG002 | CHM13-to-GRCh37 | SNP | 0.9890 | 0.9909 | 0.9900 | 3315802 | 36883 | 30406 |
| HG002 | GRCh38 | INDEL | 0.9893 | 0.9889 | 0.9891 | 519842 | 5625 | 6059 |
| HG002 | CHM13-to-GRCh38 | INDEL | 0.9895 | 0.9911 | 0.9903 | 519929 | 5538 | 4887 |
| HG002 | GRCh37 | INDEL | 0.9858 | 0.9843 | 0.9851 | 514968 | 7421 | 8551 |
| HG002 | CHM13-to-GRCh37 | INDEL | 0.9864 | 0.9903 | 0.9884 | 515300 | 7089 | 5244 |
| HG005 | GRCh38 | SNP | 0.9914 | 0.9876 | 0.9895 | 3247494 | 28120 | 40927 |
| HG005 | CHM13-to-GRCh38 | SNP | 0.9936 | 0.9903 | 0.9920 | 3254520 | 21094 | 31718 |
| HG005 | GRCh37 | SNP | 0.9861 | 0.9801 | 0.9831 | 3220745 | 45244 | 65350 |
| HG005 | CHM13-to-GRCh37 | SNP | 0.9894 | 0.9904 | 0.9899 | 3231517 | 34472 | 31338 |
| HG005 | GRCh38 | INDEL | 0.9922 | 0.9900 | 0.9911 | 413504 | 3263 | 4309 |
| HG005 | CHM13-to-GRCh38 | INDEL | 0.9923 | 0.9917 | 0.9920 | 413577 | 3190 | 3566 |
| HG005 | GRCh37 | INDEL | 0.9883 | 0.9844 | 0.9864 | 408990 | 4834 | 6685 |
| HG005 | CHM13-to-GRCh37 | INDEL | 0.9891 | 0.9912 | 0.9902 | 409333 | 4491 | 3763 |

**Table S3:** Small variant calling accuracy for 30x WGS datasets using BWA-MEM– DeepVariant in all GIAB v4.2.1 regions^22^

| Sample | Method | Type | Recall | Precision | $F_1$ | TP | FN | FP |
| --- | --- | --- | --- | --- | --- | --- | --- | --- |
| HG001 | GRCh38 | SNP | 0.9945 | 0.9983 | 0.9964 | 3236509 | 17863 | 5417 |
| HG001 | CHM13-to-GRCh38 | SNP | 0.9958 | 0.9981 | 0.9970 | 3240823 | 13549 | 6144 |
| HG001 | GRCh37 | SNP | 0.9883 | 0.9978 | 0.9931 | 3213802 | 38041 | 6937 |
| HG001 | CHM13-to-GRCh37 | SNP | 0.9908 | 0.9980 | 0.9944 | 3222057 | 29786 | 6364 |
| HG001 | GRCh38 | INDEL | 0.9925 | 0.9967 | 0.9946 | 464179 | 3514 | 1609 |
| HG001 | CHM13-to-GRCh38 | INDEL | 0.9925 | 0.9966 | 0.9946 | 464176 | 3517 | 1626 |
| HG001 | GRCh37 | INDEL | 0.9888 | 0.9966 | 0.9927 | 461638 | 5237 | 1637 |
| HG001 | CHM13-to-GRCh37 | INDEL | 0.9893 | 0.9966 | 0.9930 | 461883 | 4992 | 1623 |
| HG002 | GRCh38 | SNP | 0.9937 | 0.9990 | 0.9963 | 3343863 | 21263 | 3436 |
| HG002 | CHM13-to-GRCh38 | SNP | 0.9951 | 0.9985 | 0.9968 | 3348636 | 16490 | 4881 |
| HG002 | GRCh37 | SNP | 0.9874 | 0.9985 | 0.9929 | 3310312 | 42373 | 5059 |
| HG002 | CHM13-to-GRCh37 | SNP | 0.9898 | 0.9985 | 0.9941 | 3318525 | 34160 | 4935 |
| HG002 | GRCh38 | INDEL | 0.9919 | 0.9972 | 0.9946 | 521223 | 4244 | 1506 |
| HG002 | CHM13-to-GRCh38 | INDEL | 0.9919 | 0.9972 | 0.9945 | 521209 | 4258 | 1548 |
| HG002 | GRCh37 | INDEL | 0.9884 | 0.9971 | 0.9927 | 516310 | 6079 | 1560 |
| HG002 | CHM13-to-GRCh37 | INDEL | 0.9888 | 0.9971 | 0.9929 | 516527 | 5862 | 1561 |
| HG005 | GRCh38 | SNP | 0.9930 | 0.9986 | 0.9958 | 3252742 | 22872 | 4588 |
| HG005 | CHM13-to-GRCh38 | SNP | 0.9945 | 0.9982 | 0.9963 | 3257465 | 18149 | 5812 |
| HG005 | GRCh37 | SNP | 0.9878 | 0.9984 | 0.9931 | 3226259 | 39730 | 5227 |
| HG005 | CHM13-to-GRCh37 | SNP | 0.9904 | 0.9984 | 0.9944 | 3234785 | 31204 | 5309 |
| HG005 | GRCh38 | INDEL | 0.9929 | 0.9977 | 0.9953 | 413827 | 2941 | 993 |
| HG005 | CHM13-to-GRCh38 | INDEL | 0.9930 | 0.9976 | 0.9953 | 413845 | 2923 | 1021 |
| HG005 | GRCh37 | INDEL | 0.9892 | 0.9977 | 0.9934 | 409353 | 4472 | 969 |
| HG005 | CHM13-to-GRCh37 | INDEL | 0.9898 | 0.9977 | 0.9937 | 409620 | 4205 | 987 |

**Table S4:**
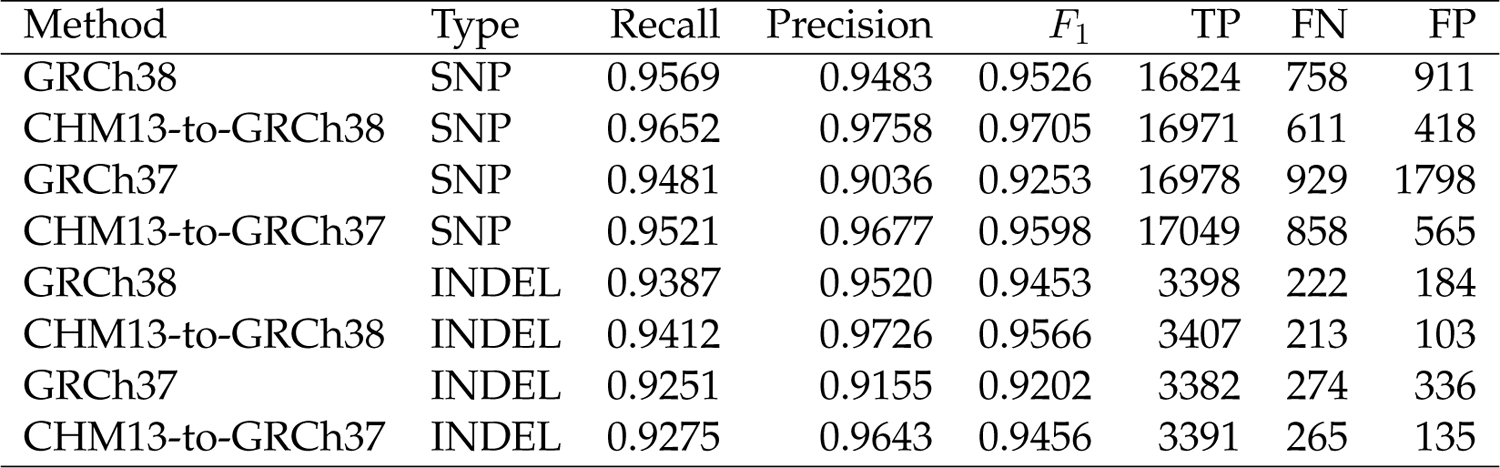
Small variant calling accuracy for 30x WGS datasets using BWA-MEM–GATK-HaplotypeCaller in GIAB CMRG regions for HG002^23^

**Table S5:** Small variant calling accuracy for 30x WGS datasets using BWA-MEM– DeepVariant in GIAB CMRG regions for HG002^23^

| Method | Type | Recall | Precision | $F_1$ | TP | FN | FP |
| --- | --- | --- | --- | --- | --- | --- | --- |
| GRCh38 | SNP | 0.9616 | 0.9854 | 0.9734 | 16907 | 675 | 250 |
| CHM13-to-GRCh38 | SNP | 0.9700 | 0.9904 | 0.9801 | 17055 | 527 | 164 |
| GRCh37 | SNP | 0.9547 | 0.9790 | 0.9667 | 17095 | 812 | 366 |
| CHM13-to-GRCh37 | SNP | 0.9591 | 0.9896 | 0.9741 | 17175 | 732 | 179 |
| GRCh38 | INDEL | 0.9340 | 0.9735 | 0.9533 | 3381 | 239 | 98 |
| CHM13-to-GRCh38 | INDEL | 0.9362 | 0.9762 | 0.9558 | 3389 | 231 | 88 |
| GRCh37 | INDEL | 0.9248 | 0.9667 | 0.9453 | 3381 | 275 | 124 |
| CHM13-to-GRCh37 | INDEL | 0.9267 | 0.9751 | 0.9503 | 3388 | 268 | 92 |

**Table S6:** Difficult regions stratified by GIAB^23^. The sizes are calculated after intersecting stratified regions with the GIAB v4.2.1 confident regions for HG002^22^

| Reference | GIAB subset name | Legend | Size |
| --- | --- | --- | --- |
| GRCh38 | alldifficultregions | AllDifficult | 415,601,891 |
| GRCh38 | gclt25orgt65_slop50 | ExtremeGC | 164,164,687 |
| GRCh38 | AllTandemRepeatsandHomopolymers_slop5 | Tandem&Homo | 121,406,218 |
| GRCh38 | lowmappabilityall | LowMap | 105,002,312 |
| GRCh38 | segdups | SegDups | 83,843,041 |
| GRCh38 | allOtherDifficultregions | OtherDifficult | 22,511,200 |
| GRCh37 | alldifficultregions | AllDifficult | 393,128,724 |
| GRCh37 | AllTandemRepeatsandHomopolymers_slop5 | Tandem&Homo | 121,049,426 |
| GRCh37 | lowmappabilityall | LowMap | 75,276,894 |
| GRCh37 | segdups | SegDups | 72,412,437 |
| GRCh37 | gclt25orgt65_slop50 | ExtremeGC | 62,521,603 |
| GRCh37 | allOtherDifficultregions | OtherDifficult | 29,884,114 |

**Table S7:** Small variant calling accuracy for 30x WGS datasets using BWA-MEM–GATK-HaplotypeCaller in GIAB difficult regions for HG002^23^

| Method | Type | Subset | Recall | Precision | $F_1$ | TP | FN | FP |
| --- | --- | --- | --- | --- | --- | --- | --- | --- |
| GRCh38 | SNP | GC<25or>65 | 0.9943 | 0.9834 | 0.9889 | 212359 | 1209 | 3576 |
| CHM13-to-GRCh38 | SNP | GC<25or>65 | 0.9954 | 0.9884 | 0.9919 | 212591 | 977 | 2487 |
| GRCh37 | SNP | GC<25or>65 | 0.9891 | 0.9759 | 0.9825 | 210064 | 2318 | 5179 |
| CHM13-to-GRCh37 | SNP | GC<25or>65 | 0.9910 | 0.9882 | 0.9896 | 210470 | 1912 | 2514 |
| GRCh38 | SNP | Tandem&Homo | 0.9948 | 0.9836 | 0.9891 | 181945 | 960 | 3059 |
| CHM13-to-GRCh38 | SNP | Tandem&Homo | 0.9951 | 0.9862 | 0.9906 | 182017 | 888 | 2564 |
| GRCh37 | SNP | Tandem&Homo | 0.9909 | 0.9795 | 0.9852 | 180733 | 1662 | 3808 |
| CHM13-to-GRCh37 | SNP | Tandem&Homo | 0.9919 | 0.9872 | 0.9896 | 180925 | 1470 | 2357 |
| GRCh38 | SNP | OtherDifficult | 0.8941 | 0.6999 | 0.7852 | 49090 | 5814 | 20974 |
| CHM13-to-GRCh38 | SNP | OtherDifficult | 0.9148 | 0.8232 | 0.8666 | 50224 | 4680 | 10750 |
| GRCh37 | SNP | OtherDifficult | 0.6308 | 0.5328 | 0.5777 | 41407 | 24230 | 36307 |
| CHM13-to-GRCh37 | SNP | OtherDifficult | 0.6828 | 0.8073 | 0.7398 | 44814 | 20823 | 10694 |
| GRCh38 | SNP | SegDups | 0.9076 | 0.8370 | 0.8709 | 109771 | 11177 | 21381 |
| CHM13-to-GRCh38 | SNP | SegDups | 0.9302 | 0.8819 | 0.9054 | 112508 | 8440 | 15071 |
| GRCh37 | SNP | SegDups | 0.8761 | 0.7342 | 0.7989 | 96344 | 13627 | 34874 |
| CHM13-to-GRCh37 | SNP | SegDups | 0.9043 | 0.8719 | 0.8878 | 99445 | 10526 | 14616 |
| GRCh38 | SNP | LowMap | 0.8806 | 0.8918 | 0.8861 | 169632 | 23008 | 20590 |
| CHM13-to-GRCh38 | SNP | LowMap | 0.9171 | 0.9241 | 0.9206 | 176665 | 15975 | 14517 |
| GRCh37 | SNP | LowMap | 0.8516 | 0.8142 | 0.8325 | 112749 | 19652 | 25729 |
| CHM13-to-GRCh37 | SNP | LowMap | 0.9006 | 0.9206 | 0.9105 | 119240 | 13161 | 10282 |
| GRCh38 | SNP | AllDifficult | 0.9611 | 0.9482 | 0.9546 | 618558 | 25007 | 33821 |
| CHM13-to-GRCh38 | SNP | AllDifficult | 0.9718 | 0.9647 | 0.9682 | 625442 | 18123 | 22947 |
| GRCh37 | SNP | AllDifficult | 0.9296 | 0.9084 | 0.9189 | 556892 | 42168 | 56269 |
| CHM13-to-GRCh37 | SNP | AllDifficult | 0.9450 | 0.9597 | 0.9523 | 566140 | 32920 | 23803 |
| GRCh38 | INDEL | GC<25or>65 | 0.9866 | 0.9869 | 0.9868 | 49835 | 675 | 680 |
| CHM13-to-GRCh38 | INDEL | GC<25or>65 | 0.9870 | 0.9894 | 0.9882 | 49854 | 656 | 549 |
| GRCh37 | INDEL | GC<25or>65 | 0.9840 | 0.9815 | 0.9828 | 49672 | 806 | 965 |
| CHM13-to-GRCh37 | INDEL | GC<25or>65 | 0.9847 | 0.9887 | 0.9867 | 49706 | 772 | 587 |
| GRCh38 | INDEL | Tandem&Homo | 0.9881 | 0.9918 | 0.9899 | 334899 | 4042 | 2941 |
| CHM13-to-GRCh38 | INDEL | Tandem&Homo | 0.9882 | 0.9923 | 0.9902 | 334933 | 4008 | 2782 |
| GRCh37 | INDEL | Tandem&Homo | 0.9861 | 0.9909 | 0.9885 | 331136 | 4665 | 3242 |
| CHM13-to-GRCh37 | INDEL | Tandem&Homo | 0.9865 | 0.9923 | 0.9894 | 331274 | 4527 | 2739 |
| GRCh38 | INDEL | OtherDifficult | 0.9573 | 0.8609 | 0.9065 | 10227 | 456 | 1762 |
| CHM13-to-GRCh38 | INDEL | OtherDifficult | 0.9591 | 0.9285 | 0.9436 | 10246 | 437 | 840 |
| GRCh37 | INDEL | OtherDifficult | 0.5957 | 0.4563 | 0.5168 | 2803 | 1902 | 3370 |
| CHM13-to-GRCh37 | INDEL | OtherDifficult | 0.6344 | 0.7358 | 0.6814 | 2985 | 1720 | 1085 |

Table S7 – *Continued from previous page*
| Method | Type | Subset | Recall | Precision | $F_1$ | TP | FN | FP |
| --- | --- | --- | --- | --- | --- | --- | --- | --- |
| GRCh38 | INDEL | SegDups | 0.9137 | 0.8599 | 0.8860 | 9888 | 934 | 1649 |
| CHM13-to-GRCh38 | INDEL | SegDups | 0.9168 | 0.9037 | 0.9102 | 9922 | 900 | 1082 |
| GRCh37 | INDEL | SegDups | 0.8876 | 0.7522 | 0.8143 | 8809 | 1115 | 2966 |
| CHM13-to-GRCh37 | INDEL | SegDups | 0.8951 | 0.8851 | 0.8901 | 8883 | 1041 | 1179 |
| GRCh38 | INDEL | LowMap | 0.8486 | 0.8678 | 0.8581 | 8836 | 1576 | 1362 |
| CHM13-to-GRCh38 | INDEL | LowMap | 0.8552 | 0.9045 | 0.8792 | 8904 | 1508 | 951 |
| GRCh37 | INDEL | LowMap | 0.8211 | 0.7691 | 0.7943 | 5917 | 1289 | 1798 |
| CHM13-to-GRCh37 | INDEL | LowMap | 0.8333 | 0.8941 | 0.8626 | 6005 | 1201 | 720 |
| GRCh38 | INDEL | AllDifficult | 0.9853 | 0.9870 | 0.9862 | 365008 | 5440 | 5096 |
| CHM13-to-GRCh38 | INDEL | AllDifficult | 0.9855 | 0.9896 | 0.9876 | 365091 | 5357 | 4051 |
| GRCh37 | INDEL | AllDifficult | 0.9808 | 0.9814 | 0.9811 | 357465 | 7014 | 7193 |
| CHM13-to-GRCh37 | INDEL | AllDifficult | 0.9816 | 0.9887 | 0.9851 | 357770 | 6709 | 4349 |

**Table S8:** Small variant calling accuracy for 30x WGS datasets using BWA-MEM– DeepVariant in GIAB difficult regions for HG002^23^

| Method | Type | Subset | Recall | Precision | $F_1$ | TP | FN | FP |
| --- | --- | --- | --- | --- | --- | --- | --- | --- |
| GRCh38 | SNP | GC<25or>65 | 0.9957 | 0.9990 | 0.9973 | 212648 | 920 | 218 |
| CHM13-to-GRCh38 | SNP | GC<25or>65 | 0.9963 | 0.9985 | 0.9974 | 212779 | 789 | 318 |
| GRCh37 | SNP | GC<25or>65 | 0.9901 | 0.9983 | 0.9942 | 210285 | 2097 | 359 |
| CHM13-to-GRCh37 | SNP | GC<25or>65 | 0.9919 | 0.9984 | 0.9951 | 210657 | 1725 | 342 |
| GRCh38 | SNP | Tandem&Homo | 0.9972 | 0.9983 | 0.9977 | 182389 | 516 | 310 |
| CHM13-to-GRCh38 | SNP | Tandem&Homo | 0.9974 | 0.9982 | 0.9978 | 182423 | 482 | 340 |
| GRCh37 | SNP | Tandem&Homo | 0.9932 | 0.9980 | 0.9956 | 181146 | 1249 | 363 |
| CHM13-to-GRCh37 | SNP | Tandem&Homo | 0.9941 | 0.9981 | 0.9961 | 181311 | 1084 | 346 |
| GRCh38 | SNP | OtherDifficult | 0.9163 | 0.9757 | 0.9450 | 50306 | 4598 | 1256 |
| CHM13-to-GRCh38 | SNP | OtherDifficult | 0.9297 | 0.9715 | 0.9502 | 51045 | 3859 | 1497 |
| GRCh37 | SNP | OtherDifficult | 0.6414 | 0.9530 | 0.7668 | 42101 | 23536 | 2076 |
| CHM13-to-GRCh37 | SNP | OtherDifficult | 0.6901 | 0.9685 | 0.8059 | 45294 | 20343 | 1474 |
| GRCh38 | SNP | SegDups | 0.9169 | 0.9847 | 0.9496 | 110896 | 10052 | 1729 |
| CHM13-to-GRCh38 | SNP | SegDups | 0.9332 | 0.9731 | 0.9528 | 112872 | 8076 | 3117 |
| GRCh37 | SNP | SegDups | 0.8847 | 0.9768 | 0.9284 | 97286 | 12685 | 2316 |
| CHM13-to-GRCh37 | SNP | SegDups | 0.9082 | 0.9752 | 0.9405 | 99877 | 10094 | 2537 |
| GRCh38 | SNP | LowMap | 0.9014 | 0.9851 | 0.9414 | 173653 | 18987 | 2634 |
| CHM13-to-GRCh38 | SNP | LowMap | 0.9275 | 0.9779 | 0.9520 | 178675 | 13965 | 4038 |
| GRCh37 | SNP | LowMap | 0.8773 | 0.9772 | 0.9246 | 116156 | 16245 | 2705 |
| CHM13-to-GRCh37 | SNP | LowMap | 0.9136 | 0.9752 | 0.9434 | 120959 | 11442 | 3077 |
| GRCh38 | SNP | AllDifficult | 0.9685 | 0.9951 | 0.9816 | 623274 | 20291 | 3073 |
| CHM13-to-GRCh38 | SNP | AllDifficult | 0.9758 | 0.9929 | 0.9842 | 627965 | 15600 | 4517 |
| GRCh37 | SNP | AllDifficult | 0.9366 | 0.9920 | 0.9635 | 561083 | 37977 | 4534 |
| CHM13-to-GRCh37 | SNP | AllDifficult | 0.9489 | 0.9922 | 0.9701 | 568432 | 30628 | 4467 |
| GRCh38 | INDEL | GC<25or>65 | 0.9895 | 0.9959 | 0.9927 | 49979 | 531 | 211 |
| CHM13-to-GRCh38 | INDEL | GC<25or>65 | 0.9895 | 0.9960 | 0.9927 | 49978 | 532 | 205 |
| GRCh37 | INDEL | GC<25or>65 | 0.9869 | 0.9958 | 0.9913 | 49815 | 663 | 216 |
| CHM13-to-GRCh37 | INDEL | GC<25or>65 | 0.9871 | 0.9959 | 0.9915 | 49826 | 652 | 213 |
| GRCh38 | INDEL | Tandem&Homo | 0.9914 | 0.9962 | 0.9938 | 336023 | 2918 | 1360 |
| CHM13-to-GRCh38 | INDEL | Tandem&Homo | 0.9913 | 0.9961 | 0.9937 | 336001 | 2940 | 1385 |
| GRCh37 | INDEL | Tandem&Homo | 0.9894 | 0.9962 | 0.9928 | 332240 | 3561 | 1346 |
| CHM13-to-GRCh37 | INDEL | Tandem&Homo | 0.9896 | 0.9962 | 0.9928 | 332294 | 3507 | 1366 |
| GRCh38 | INDEL | OtherDifficult | 0.9639 | 0.9899 | 0.9767 | 10299 | 386 | 110 |
| CHM13-to-GRCh38 | INDEL | OtherDifficult | 0.9622 | 0.9897 | 0.9758 | 10281 | 404 | 112 |
| GRCh37 | INDEL | OtherDifficult | 0.6066 | 0.9634 | 0.7445 | 2854 | 1851 | 109 |
| CHM13-to-GRCh37 | INDEL | OtherDifficult | 0.6391 | 0.9737 | 0.7717 | 3007 | 1698 | 82 |

Table S8 – *Continued from previous page*
| Method | Type | Subset | Recall | Precision | $F_1$ | TP | FN | FP |
| --- | --- | --- | --- | --- | --- | --- | --- | --- |
| GRCh38 | INDEL | SegDups | 0.9236 | 0.9874 | 0.9544 | 9995 | 827 | 130 |
| CHM13-to-GRCh38 | INDEL | SegDups | 0.9250 | 0.9859 | 0.9545 | 10010 | 812 | 146 |
| GRCh37 | INDEL | SegDups | 0.8972 | 0.9845 | 0.9388 | 8904 | 1020 | 143 |
| CHM13-to-GRCh37 | INDEL | SegDups | 0.9029 | 0.9854 | 0.9423 | 8960 | 964 | 135 |
| GRCh38 | INDEL | LowMap | 0.8681 | 0.9825 | 0.9218 | 9040 | 1373 | 163 |
| CHM13-to-GRCh38 | INDEL | LowMap | 0.8720 | 0.9805 | 0.9231 | 9080 | 1333 | 182 |
| GRCh37 | INDEL | LowMap | 0.8425 | 0.9782 | 0.9053 | 6073 | 1135 | 137 |
| CHM13-to-GRCh37 | INDEL | LowMap | 0.8511 | 0.9796 | 0.9109 | 6134 | 1073 | 129 |
| GRCh38 | INDEL | AllDifficult | 0.9889 | 0.9962 | 0.9925 | 366326 | 4122 | 1470 |
| CHM13-to-GRCh38 | INDEL | AllDifficult | 0.9888 | 0.9961 | 0.9925 | 366308 | 4140 | 1509 |
| GRCh37 | INDEL | AllDifficult | 0.9842 | 0.9960 | 0.9901 | 358734 | 5746 | 1517 |
| CHM13-to-GRCh37 | INDEL | AllDifficult | 0.9848 | 0.9960 | 0.9904 | 358931 | 5549 | 1519 |

**Table S9:** Small variant calling accuracy for 28x WGS PacBio-HiFi data using minimap2– DeepVariant in all GIAB v4.2.1 regions for HG002^22^

| Method | Type | Recall | Precision | $F_1$ | TP | FN | FP |
| --- | --- | --- | --- | --- | --- | --- | --- |
| GRCh38 | SNP | 0.9988 | 0.9991 | 0.9990 | 3361165 | 3961 | 3012 |
| CHM13-to-GRCh38 | SNP | 0.9988 | 0.9990 | 0.9989 | 3361055 | 4071 | 3482 |
| GRCh37 | SNP | 0.9923 | 0.9979 | 0.9951 | 3326880 | 25805 | 7129 |
| CHM13-to-GRCh37 | SNP | 0.9945 | 0.9986 | 0.9965 | 3334094 | 18591 | 4667 |
| GRCh38 | INDEL | 0.9467 | 0.9190 | 0.9327 | 497477 | 27990 | 45387 |
| CHM13-to-GRCh38 | INDEL | 0.9466 | 0.9189 | 0.9326 | 497426 | 28041 | 45429 |
| GRCh37 | INDEL | 0.9436 | 0.9184 | 0.9308 | 492918 | 29472 | 45365 |
| CHM13-to-GRCh37 | INDEL | 0.9448 | 0.9187 | 0.9315 | 493541 | 28849 | 45229 |

**Table S10:** Small variant calling accuracy for 28x WGS PacBio-HiFi data using minimap2–DeepVariant in GIAB CMRG regions for HG002^23^

| Method | Type | Recall | Precision | $F_1$ | TP | FN | FP |
| --- | --- | --- | --- | --- | --- | --- | --- |
| GRCh38 | SNP | 0.9876 | 0.9795 | 0.9836 | 17364 | 218 | 365 |
| CHM13-to-GRCh38 | SNP | 0.9952 | 0.9890 | 0.9921 | 17497 | 85 | 196 |
| GRCh37 | SNP | 0.9867 | 0.9698 | 0.9782 | 17668 | 239 | 553 |
| CHM13-to-GRCh37 | SNP | 0.9870 | 0.9702 | 0.9785 | 17675 | 232 | 547 |
| GRCh38 | INDEL | 0.8851 | 0.8700 | 0.8775 | 3204 | 416 | 496 |
| CHM13-to-GRCh38 | INDEL | 0.8876 | 0.8760 | 0.8817 | 3213 | 407 | 470 |
| GRCh37 | INDEL | 0.8816 | 0.8690 | 0.8753 | 3223 | 433 | 503 |
| CHM13-to-GRCh37 | INDEL | 0.8829 | 0.8716 | 0.8772 | 3228 | 428 | 491 |

**Table S11:**
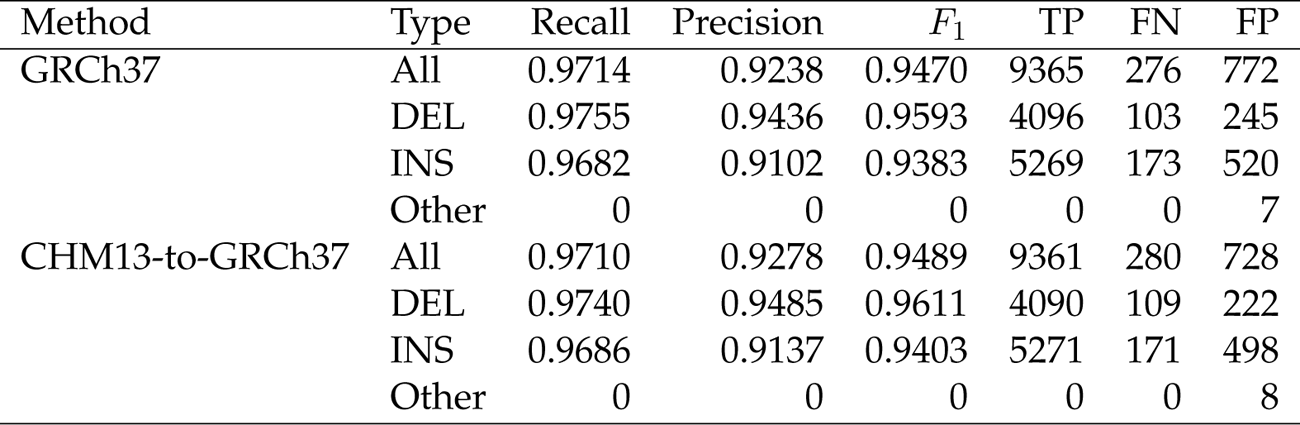
Structural variant calling accuracy for 28x WGS PacBio-HiFi data using minimap2–Sniffles 2 in GRCh37 GIAB Tier 1 benchmark regions for HG002^24^

| Method | Type | Recall | Precision | $F_1$ | TP | FN | FP |
| --- | --- | --- | --- | --- | --- | --- | --- |
| GRCh37 | All | 0.9714 | 0.9238 | 0.9470 | 9365 | 276 | 772 |
|  | DEL | 0.9755 | 0.9436 | 0.9593 | 4096 | 103 | 245 |
|  | INS | 0.9682 | 0.9102 | 0.9383 | 5269 | 173 | 520 |
|  | Other | 0 | 0 | 0 | 0 | 0 | 7 |
| CHM13-to-GRCh37 | All | 0.9710 | 0.9278 | 0.9489 | 9361 | 280 | 728 |
|  | DEL | 0.9740 | 0.9485 | 0.9611 | 4090 | 109 | 222 |
|  | INS | 0.9686 | 0.9137 | 0.9403 | 5271 | 171 | 498 |
|  | Other | 0 | 0 | 0 | 0 | 0 | 8 |

**Table S12:** Structural variant calling accuracy for 25x WGS PacBio-HiFi data using minimap2–Sniffles 2 in GRCh38 GIAB CMRG regions for HG002^23^

| Method | Type | Recall | Precision | $F_1$ | TP | FN | FP |
| --- | --- | --- | --- | --- | --- | --- | --- |
| GRCh38 | All | 0.9677 | 0.9519 | 0.9598 | 198 | 7 | 10 |
|  | DEL | 0.9785 | 0.9381 | 0.9579 | 91 | 2 | 6 |
|  | INS | 0.9609 | 0.9685 | 0.9647 | 123 | 5 | 4 |
| CHM13-to-GRCh38 | All | 0.9677 | 0.9612 | 0.9644 | 198 | 7 | 8 |
|  | DEL | 0.9785 | 0.9579 | 0.9681 | 91 | 2 | 4 |
|  | INS | 0.9609 | 0.9685 | 0.9647 | 123 | 5 | 4 |

